# Elementary vectors reveal minimal interactions in microbial communities

**DOI:** 10.1101/2025.07.30.667663

**Authors:** Stefan Müller, Michael Predl, Diana Széliová, Jürgen Zanghellini

**Affiliations:** Faculty of Mathematics, University of Vienna, Oskar Morgenstern Platz 1, 1090, Vienna, Austria; Department of Analytical Chemistry, University of Vienna, Währinger Strasse 42, 1090, Vienna, Austria; Research Network Data Science, University of Vienna, Kolingasse 14-16, 1090, Vienna, Austria

**Keywords:** microbial communities, metabolic modeling, polyhedral geometry

## Abstract

Understanding microbial communities is essential for progress in ecology, biotechnology, and human health. In the last decade, constraint-based metabolic models of individual organisms have been combined to study microbial consortia. In this work, we present a geometric framework for characterizing all feasible microbial interactions. We project community models onto the key variables of interaction: exchange fluxes and community compositions. Based on this projection, we compute elementary composition/exchange fluxes (ECXs), extending the concept of minimal metabolic pathways from individual species to entire communities. Every feasible metabolic state of a community can be expressed as a combination of these elementary vectors. Notably, each ECX corresponds to a distinct ecological interaction type, such as specialization, commensalism, or mutualism. Finally, our geometric formulation enables the direct application of existing constraint-based methods, such as flux variability analysis and minimal cut sets, to microbial communities, providing a foundation for rational community design.

## Introduction

Microbial communities play a key role in nature. They are essential for human health [18, 9], support global geochemical cycles [22, 36], and have various biotechnological applications such as the production of valuable compounds or wastewater treatment [7, 25].

Within microbial communities, members engage in complex interactions involving nutrient exchange, leading to various relationships such as mutualism, commensalism, and specialization [31]. Studying these interactions in laboratory settings, however, remains challenging. Metaomics approaches—including genomics, transcriptomics, and metabolomics—allow for extensive characterization of microbial communities [23], but fall short in revealing specific interactions among community members or how community composition and growth rate depend on environmental factors. Nevertheless, these datasets lay the groundwork for metabolic modeling, which offers a promising approach to address these questions [45].

Single-species models capture a cell’s metabolic capabilities and its interactions with the environment. By extending this approach to microbial communities, one aims to map the range of potential feeding interactions, community compositions, and growth rates, and to assess the robustness of these communities against environmental perturbations [12, 45].

Constraint-based metabolic modeling (CBM) offers a promising approach for studying microbial communities. A key assumption in CBM is metabolic steady state. In community models, this implies balanced growth, where all species grow at the same exponential rate to maintain a stable composition. While individual growth rates may fluctuate over time, the community members must grow at the same rate on average; otherwise, the fastest-growing member would outcompete the others. This balanced growth assumption underlies several community modeling approaches such as cFBA [26], OptDeg [29], SteadyCom [8], and RedCom [30]. Experimental evidence also supports balanced growth; for example, observations suggest that the human gut microbiome can maintain a stable average composition over extended periods, lasting months or even years [16, 10].

Extending CBM methods from the single-species to the community level introduces significant technical challenges. Community models are inherently nonlinear, involving either products of fluxes and compositions or, after reformulation, products of growth rate and compositions [24]. Although nonlinear optimization techniques can in principle be applied, they become computationally infeasible when many species or large-scale metabolic networks are considered. A common approach is to linearize the problem by fixing either compositions [26, 29] or growth rate [8, 30], thereby enabling the use of established CBM methods. For instance, single-species models can be reduced by selecting elementary flux vectors with minimal conversions [30].

In this work, we scale reaction rates by community, rather than single-species biomass. For fixed growth rate, the resulting model is linear in both fluxes and community compositions. Rather than simplifying single-species models, we treat the full community model as an *inequality system, project* it onto community compositions and exchange fluxes, and introduce corresponding *elementary vectors*. In this way, we extend elementary pathway analysis from the single-species to the community level. The resulting *elementary compositions/exchange fluxes* represent “minimal” communities together with the interactions between their members. Crucially, these elementary vectors can be interpreted as fundamental ecological interactions such as mutualism, commensalism, or specialization.

### Organization of the work

We first recall dynamic models for single-species growth and exchange with the medium. After assuming steady state, we consider constraint-based models. As our new contribution, we formulate several versions of a community model (in terms of species-scale, community-scale, and projected fluxes, respectively) and analyze their geometry. Notably, we introduce elementary vectors for community models and state corresponding decomposition results. We illustrate all concepts and results in a running example (of a two-species community) and discuss their ecological significance.

#### Notation

We denote the real numbers by ℝ. For a finite index set Ind, we write ***x*** ∈ ℝ^Ind^ for the vector (formal sum) ∑_*i*∈Ind_ *x*_*i*_ *i* with *x*_*i*_ ∈ ℝ. (In the standard case, where Ind = {1, …, *n*}, we write ***x*** = (*x*_1_, …, *x*_*n*_)^⊤^ ∈ ℝ^*n*^.) For two vectors ***x, y*** ∈ ℝ^Ind^, we denote their scalar product by ***x*** · ***y*** = ∑_*i*∈Ind_ *x*_*i*_*y*_*i*_. We use bold letters for matrices and vectors.

## Towards constraint-based community models

We consider a set Spc = {1, …, *s*} of *s* microbial species, see the schematic diagram in Fig. 1. In particular, we denote the (intracellular) stoichiometric matrix of species *i* ∈ Spc by

**Fig. 1.**
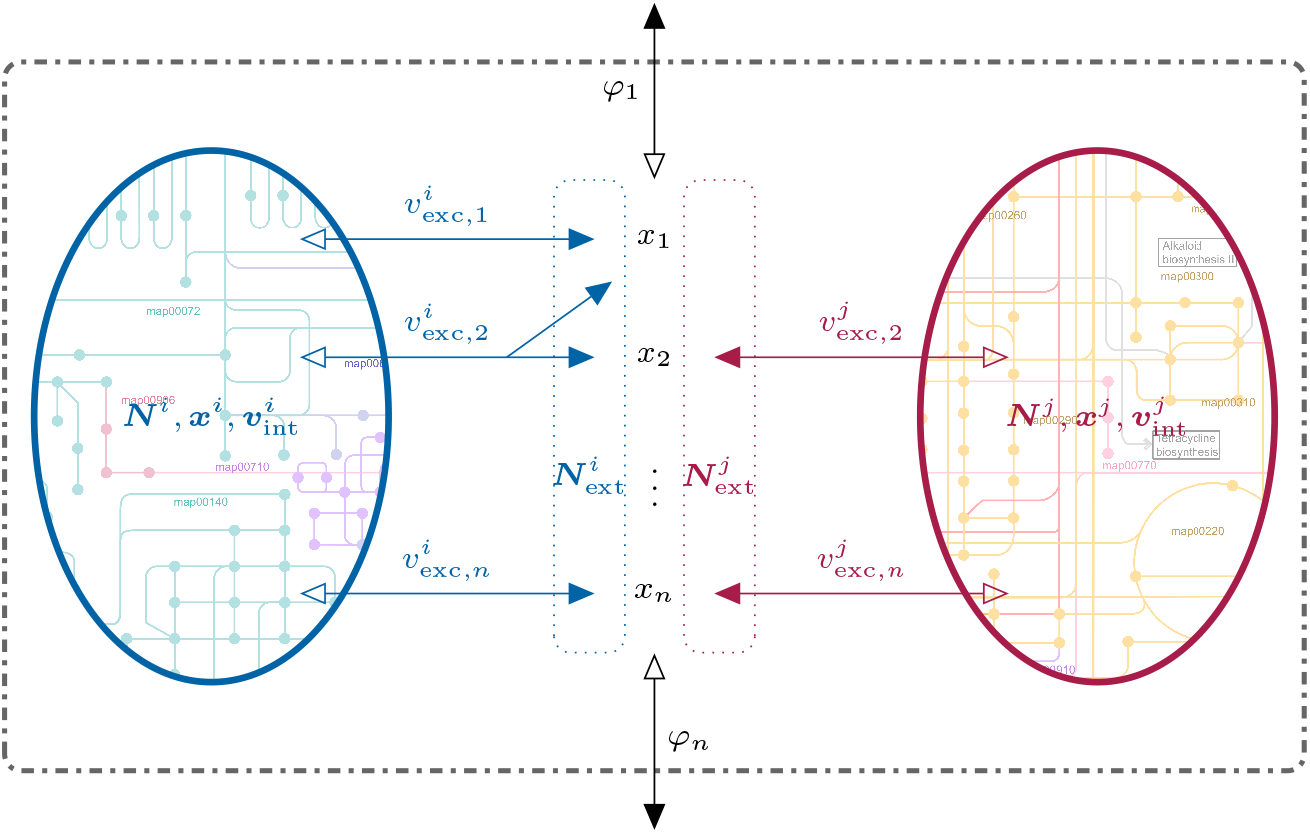
Microbial community involving species *i* and *j*. The metabolism of the corresponding cells (blue and red ovals) is given by the intracellular stoichiometries/concentrations/fluxes ***N***^*i*^, ***x***^*i*^, ***v***^*i*^ and ***N***^*j*^, ***x***^*j*^, ***v***^*j*^. The transport of the extracellular metabolites (indicated by their concentrations ***x***) across the cell membranes occurs via the exchange fluxes 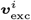 and 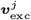 (blue and red arrows). The effect of the exchange reactions on the medium is specified by the extracellular stoichiometric matrices 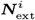 and 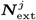 (blue and red dotted rectangles). Extracellular metabolites may have in/outflows ***φ*** to/from the medium. (Fluxes are counted positive in the direction of full arrowheads.)

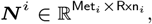

where Met_*i*_ is the set of intracellular metabolites of species *i* and Rxn_*i*_ is the set of (exchange and internal) reactions. That is, Rxn_*i*_ = Rxn_exc,*i*_ ∪ Rxn_int,*i*_ and 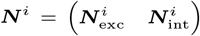. Note that ***N***^*i*^ does not involve the “biomass reaction”, used to model cellular growth in CBMs such as flux balance analysis (FBA) and elementary flux mode (EFM) analysis.

### Single species growth

Let 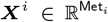 denote the amounts of substance of the free metabolites of species *i*. Further, since metabolites can be bound within macromolecules (enzymes, other proteins, DNA, RNA, etc.), let 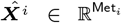 denote the total amounts of substance of the (free and bound) metabolites. Finally, let 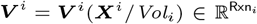 denote the reaction rates, depending on the volume-specific concentrations. Since we do not consider kinetic, but constraint-based models in the end, we do not scale by volume. Indeed, we scale by biomass *M*_*i*_,

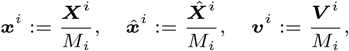

introduce growth rate^1^

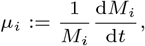

and obtain the “dynamic constraint”

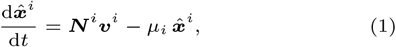

using next-generation metabolic models (explicitly including macromolecules) [40]. Note that in the derivation of FBA models, one assumes steady state and often neglects free metabolites [11].

The “dilution by growth” term 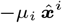 is viewed as the effect of a *biomass reaction*. To obtain the correct dimensions (a dimensionless *stoichiometric vector* times a reaction rate), we use 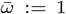 g/mmol to define the stoichiometric vector 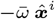 and the rate 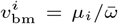 of the biomass reaction. Explicitly, the biomass reaction can be written as

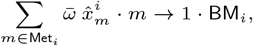

involving the metabolites *m* ∈ Met_*i*_ and the biomass “species” BM_*i*_, having molar mass 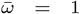 g/mmol. That is, the (dimensionless) stoichiometric coefficients 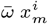, for *m* ∈ Met_*i*_, and 1 represent the synthesis of one biomass “molecule”.^2^

With molar masses 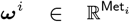 of the intracellular metabolites, the definition of biomass

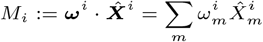

leads to the constraint

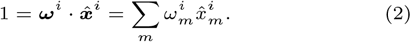

The dynamical system (1) and the mass constraint (2) allow to express growth rate in terms of reaction rates,

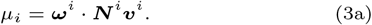

Finally, we introduce the extracellular stoichiometric matrix

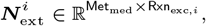

where Met_med_ is the set of *all* metabolites in the medium (even if they are not exchanged with species *i*). Now, we can define a “total” stoichiometric matrix

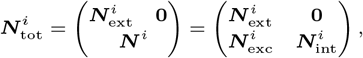

where 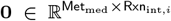 is the zero matrix, and write mass conservation as

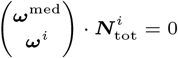

with molar masses 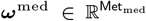 of the metabolites in the medium. As a result, we obtain growth rate in terms of *exchange* reaction rates only,

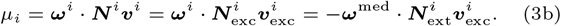

### Exchange with medium

Let 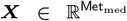 denote the amounts of substance of the metabolites in the medium. Then,

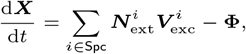

that is, the dynamics is determined by a sum involving the exchange fluxes of all microbial species and the *net* out/inflows **Φ** (from/to the medium).

To highlight the contributions of the individual species, we scale their biomass by total biomass,

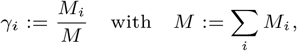

and obtain

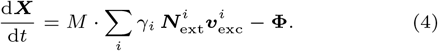

Clearly, the microbial mass fractions ***γ*** ∈ ℝ^*s*^ satisfy

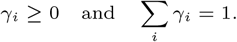

They are also called (relative) abundances or (community) composition. Further, the growth rate of total biomass

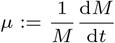

fulfills

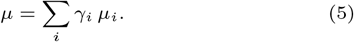

Eqn. (4), multiplied with the molar masses of the metabolites in the medium, Eqn. (3b), expressing growth rate as a function of exchange reaction rates, and Eqn. (5) relate growth rate and net out/inflows,

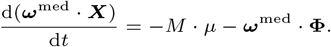

Alternatively, we can write Eqn. (4) in terms of more standard, scaled quantities. Assuming a fixed volume of the medium, *V*_med_, we introduce the concentrations ***x*** := ***X****/V*_med_, the biomass density *ρ* := *M/V*_med_, and ***φ*** := **Φ***/V*_med_, and obtain

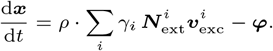

### From dynamics to steady state

In the growth model Eqn. (1) of species *i* ∈ Spc, we assume steady state or “balanced growth”. That is, metabolite concentrations and growth rate are constant over time, and we obtain

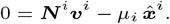

In particular, balanced growth implies that all microbial species (that are present) grow at the same rate: In the setting of a batch reactor, let *µ*_max_ := max_*i*_ *µ*_*i*_. By the definitions of ***µ*** and ***γ***, if *µ*_*i*_ *< µ*_max_ for some *i* ∈ Spc, then *γ*_*i*_ → 0 as *t* → ∞. That is, *µ*_*i*_ = *µ* for all *i* ∈ Spc (with *γ*_*i*_ *>* 0). In the setting of a flow reactor, all growth rates converge to dilution rate, *µ*_*i*_ → *d* as *t* → ∞. Again, *µ*_*i*_ = *µ*.

After extending the stoichiometric matrix by the (stoichiometric vector of the) biomass reaction,

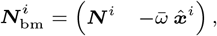

we can write steady state in “standard form”,

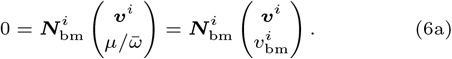

In the absence of kinetic information, we further consider lower and upper bounds 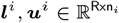 potentially for all fluxes,^3^

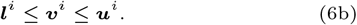

Altogether, the set of feasible growth rates *µ*_*i*_ and fluxes 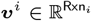 is given by the linear equation/inequality system (6). For variable *µ*_*i*_, Eqn. (6a) is homogeneous in (***v***^*i*^, *µ*_*i*_); for fixed *µ*_*i*_, it is inhomogeneous. In any case, the full system (6) is inhomogeneous.

One can further study the single-species models (6) individually, in particular, one can compute their elementary conversion vectors (thereby extending the notion of elementary conversion modes from cones to polyhedra). However, in this work, we first construct community models as *inequality systems* and analyze them via *elementary vectors* in a second step.

In principle, the (scaled) *dynamic* model (4) for the medium can be combined with the constraint-based models (6) for the individual species, as in dynamic FBA. However, in this work, we consider also Eqn. (4) at steady state,

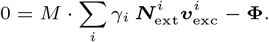

Let *j* ∈ Met_med_ be a metabolite in the medium and assume that Φ_*j*_ = 0, that is, there is no net out/inflow of this metabolite (from/to the medium). Then,

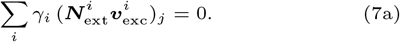

If one assumes either net inflow (Φ_*j*_ ≤ 0) or net outflow (Φ_*j*_ ≥ 0), then either

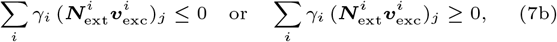

respectively. If both in- and outflow are allowed, then we do not obtain a constraint (for metabolite *j*). Hence, we introduce three subsets of Met_med_, corresponding to no in/outflow, net inflow, and net outflow, denoted by 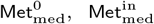, and 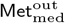 respectively.^4^

In the next section, we formulate several versions of constraint-based community models. Similarly to [30], we introduce systems (8) and (9), expressed in terms of species-scale and community-scale fluxes, respectively. We show that the two formulations are equivalent and analyze their geometry. Most importantly, we project system (9) onto the variables of interest, exchange fluxes and microbial mass fractions, to obtain system (11), and we define the combined composition/exchange cone and polytope in Eqns. (12).

## Results: community geometry

As a first version of a constraint-based community model, we combine Eqns. (6) and (7) and obtain the following system for growth rate *µ*, microbial mass fractions ***γ*** ∈ ℝ^*s*^, and fluxes 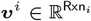, for *i* ∈ Spc, see also [30].

Community model in *µ*, ***γ*** and ***v***^*i*^:

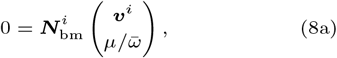

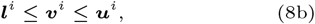

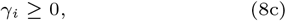

for *i* ∈ Spc, and

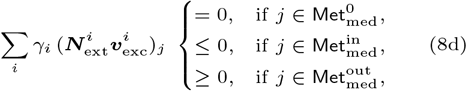

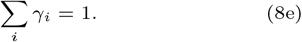

Note that Eqn. (8d) is bilinear in ***γ*** and ***v***^*i*^, *i* ∈ Spc. As in [30], we multiply Eqns. (8ab) with *γ*_*i*_ and perform a change of variables. Instead of (*µ*, ***γ, v***^1^, …, ***v***^*s*^), we consider 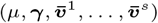, where

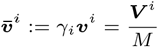

are rates/fluxes scaled by total biomass (rather than by individual biomasses).

As it turns out, an equivalent version of a constraint-based community model is given by the following system for growth rate *µ*, microbial mass fractions ***γ*** ∈ ℝ^*s*^, and (scaled) fluxes 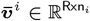, *i* ∈ Spc. (For a glossary of notation, see Table 3.)

Community model in *µ*, ***γ*** and 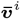:

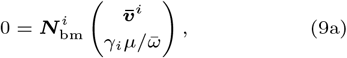

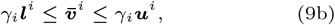

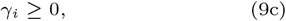

for *i* ∈ Spc, and

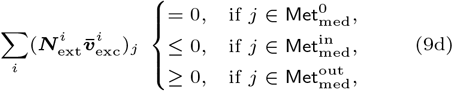

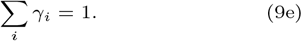

Systems (8) and (9) are equivalent only since they involve the constraints (8b) and (9b), respectively. Just assume that species *i* is not present in the community, that is, *γ*_*i*_ = 0. On the one hand, (8ab) specifies constraints for the fluxes ***v***^*i*^, although they do not contribute to (8d). On the other hand, the fluxes 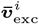 do contribute to (9d), and only (9b) ensures 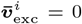. That is the reason why we include lower and upper bounds for all exchange fluxes. (In fact, bounds are required at least for the uptake fluxes of all metabolites with net inflow to the medium.)

For fixed *µ*, system (9) is linear, in particular, homogeneous in the remaining variables 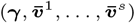, except for constraint (9e), the normalization of mass fractions. In particular, the single-species submodel (9abc) is the homogenization of system (8ab), with homogenization variable *γ*_*i*_.

In the following, we often drop (9e) for the moment, and reconsider it at the end of our analysis. Indeed, whereas system (9) defines a polytope in 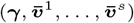, system (9abcd) defines an (essentially isomorphic) polyhedral cone, for which projection methods are readily available.

**Note** (computation). Since system (9) is linear for fixed *µ* (or fixed ***γ***), it is accessible to standard CBM approaches like FBA, flux variability analysis (FVA), and EFM analysis. In particular, for fixed *µ*, the mass fractions *γ*_*i*_ can be interpreted as fluxes. This translation can be conveniently performed automatically using PyCoMo [44], a Python package for generating (genome-scale) community models compatible with current openCOBRA file formats.

### Projecting to mass fractions and/or exchange fluxes

Ultimately, we are interested in a community model that involves only microbial mass fractions and/or exchange fluxes. In the following, we eliminate internal fluxes, which only appear in the single-species subsystems (9abc).

For species *i* and fixed growth rate *µ*, the set of feasible microbial mass fractions *γ*_*i*_ ∈ ℝ and feasible fluxes 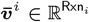 is given by the homogeneous equation/inequality system (9abc), defining the polyhedral cone

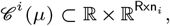

depending on growth rate. We denote its projection to *γ*_*i*_ ∈ ℝ and the exchange fluxes 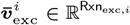 by

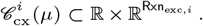

Explicitly,

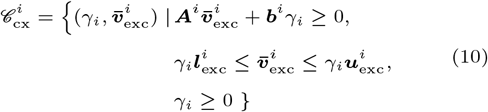

for some matrix 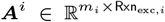 and some vector 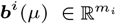, depending linearly on *µ* (and on 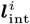 and 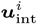). Of course, the latter two inequalities can be included in the first one. For clarity, we explicitly restate the inequalities that did not involve internal fluxes.

Using the explicit form (10), we obtain the desired version of a constraint-based community model, involving growth rate *µ*, microbial mass fractions ***γ*** ∈ ℝ^*s*^, and exchange fluxes 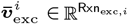, *i* ∈ Spc.

Projected community model in *µ*, ***γ*** and 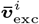:

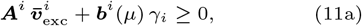

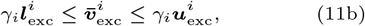

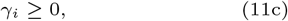

for *i* ∈ Spc, and

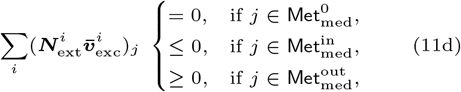

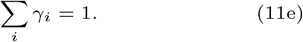

The projections (11abc) of the single-species subsystems (9abc) could be computed in a preprocessing step. For clarity in the derivation of the community model, we first considered the full single-species subsystems (8abc) and (9abc) and eliminated internal fluxes in a subsequent step.

**Note** (computation). The basic method to eliminate variables from a system of linear inequalities is the Fourier-Motzkin algorithm [20, 21, 37], see also [46, 33]. For the examples in this work, we use block elimination [4]. Essentially, the new inequalities (for the remaining variables) are obtained by forming nonnegative linear combinations of the original inequalities. For a previous use of this method in the analysis of metabolic networks, see [32].

For fixed growth rate *µ*, system (11) defines the (combined) *composition/exchange polytope*

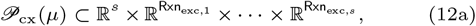

and the homogeneous part (11abcd) defines the (combined) *composition/exchange cone*

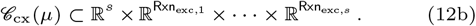

Obviously, 𝒞_cx_ is a polyhedral cone, and 𝒫_cx_ is a bounded polyhedron (that is, a polytope). In fact, we can say more about the two objects.

#### Proposition 1.

𝒞_cx_ *is pointed, and* 𝒞_cx_ *and* 𝒫_cx_ *are isomorphic in the sense that rays of* 𝒞_cx_ *are in one-to-one correspondence with points of* 𝒫_cx_.

*Proof* First, we show that the cone 𝒞_cx_ is pointed. Assume that both 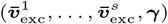 and 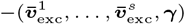 are elements of 𝒞_cx_. Then, ***γ*** = 0 by Eqn. (11c) which further implies 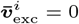 for *i* ∈ Spc by Eqn. (11b).

Second, we exhibit the isomorphism. On the one hand, for every point 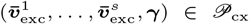 and every *λ >* 0, we find 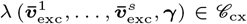 (since Eqn. (11e) is discarded, when going from 𝒫_cx_ to 𝒞_cx_), that is, the corresponding ray is contained in the cone. On the other hand, every ray of the cone is given by some nonzero point 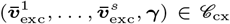 with nonzero ***γ***. Hence, there exists a unique *λ >* 0 such that 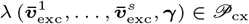, in particular, *λ* ∑ *γ*_*i*_ = 1. □

By further projecting 𝒫_cx_ and 𝒞_cx_ to the microbial mass fractions ***γ*** ∈ ℝ^*s*^, we obtain the *composition polytope* and *composition cone*

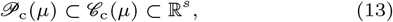

respectively. Clearly, 𝒞_c_ is pointed, and 𝒞_c_ and 𝒫_c_ are isomorphic, as above.

Alternatively, we can eliminate the microbial mass fractions. By projecting 𝒫_cx_ and 𝒞_cx_ to the exchange fluxes 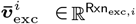, *i* ∈ Spc, we obtain the *exchange polytope* and *exchange cone*

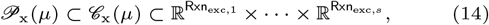

respectively. On the one hand, this projection assigns to every point in 𝒫_cx_ a unique point in 𝒫_x_. On the other hand, Eqn. (3b) implies

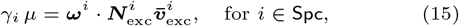

which assigns to every point in 𝒫_x_ a *unique* point in 𝒫_cx_, provided that *µ >* 0. In this case, 𝒫_cx_ is just the image of 𝒫_x_ under a linear transformation. We have shown:

#### Proposition 2.

𝒫_x_ *is in one-to-one correspondence with* 𝒫_cx_.

As above, 𝒞_x_ is pointed, and 𝒞_x_ and 𝒫_x_ are isomorphic.

For an overview of all geometric objects introduced above, see Table 1.

**Table 1.**
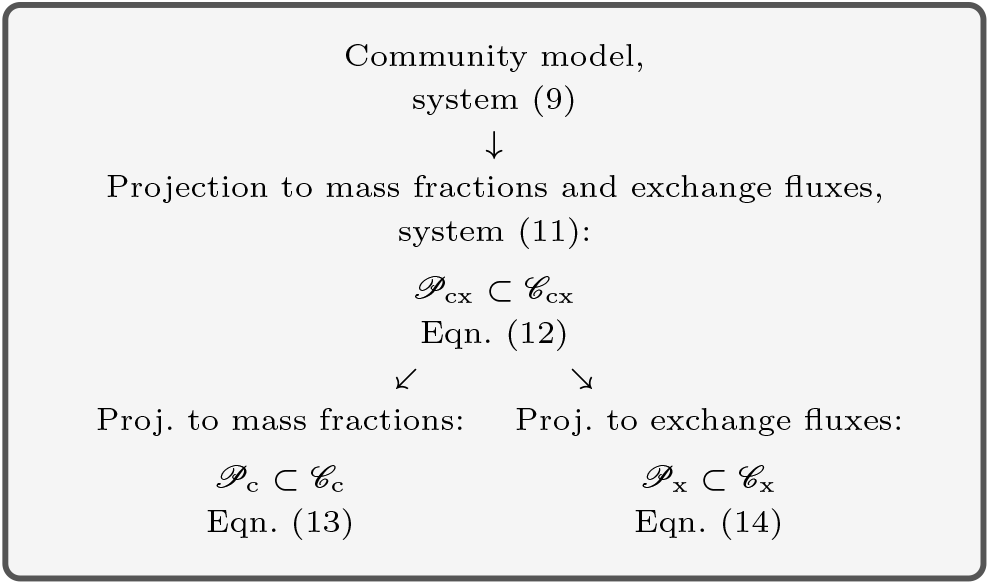
The hierarchy of (projected) community models and the related polytopes and polyhedral cones.

**Example 1**. We consider a microbial community involving just two species, as shown in Fig. 2a. There is an inflow of metabolites *X*^1^ and *X*^2^ to the medium and an outflow of *Y*^1^ and *Y*^2^, that is, 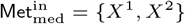 and 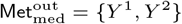.

**Fig. 2.**
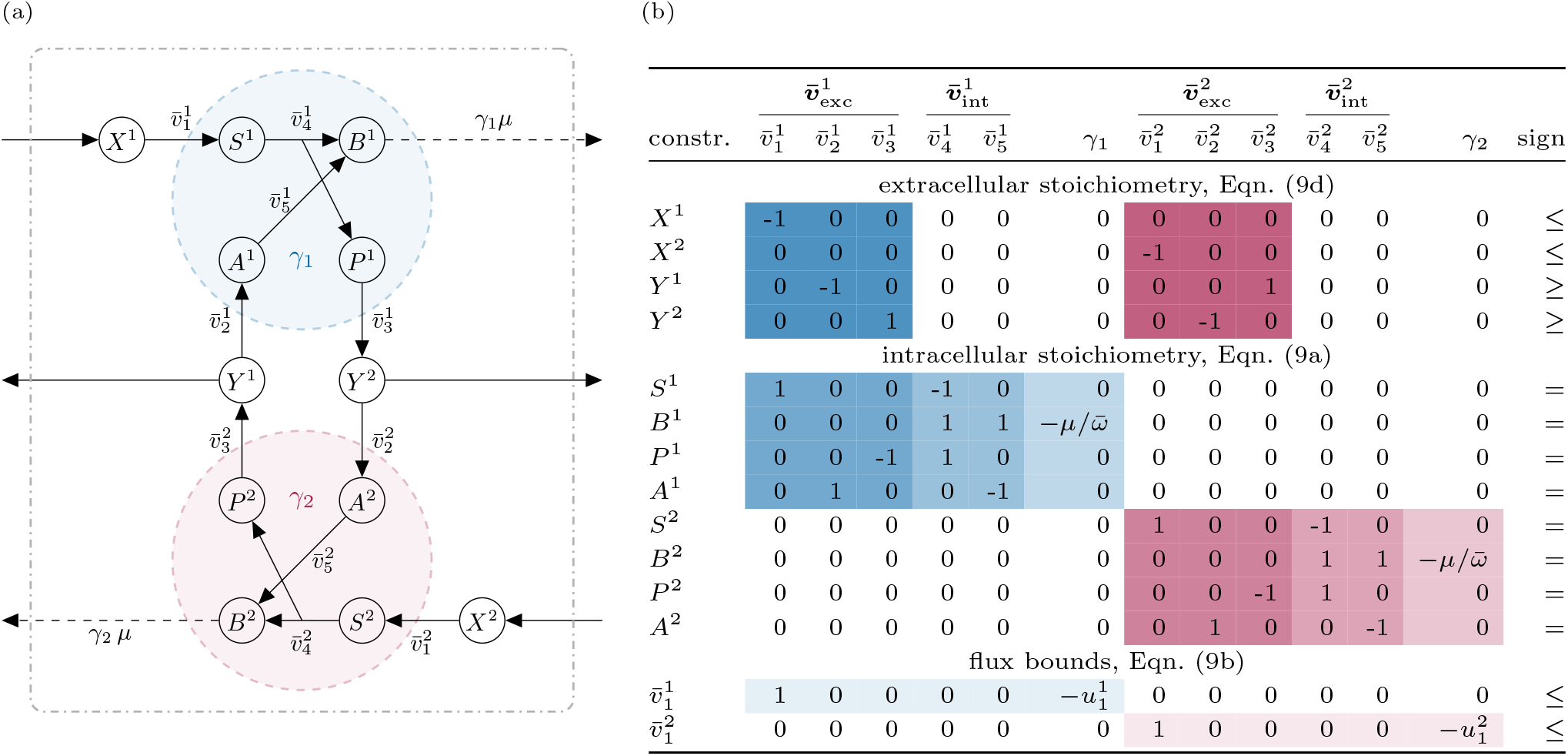
Two-species community: (a) reaction network with all (internal and exchange) fluxes and all in/outflows to/from the medium. (b) matrix representation of the (in-)equality constraints (9abd) for the exchange fluxes 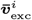, the internal fluxes 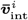, and the mass fractions *γ*_*i*_ (of the two species). The constraints arise from the steady-state assumptions (9d) and (9a) for the extra- and intracellular metabolites as well as from the flux bounds (9b). The sign of each (in)equality is given by the last column. Blue and red colors indicate sub-matrices corresponding to species one and two, respectively. Within each color, progressively lighter shadings denote the sub-matrices 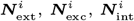, and 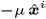 (the “dilution by growth” term). In particular, constraints (9d) involve the extracellular matrices 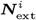 (darkest shading), whereas constraints (9a) can be expanded into 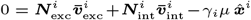, using 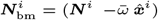 and 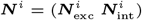, and hence involve the intracellular matrices 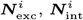, and the vector 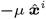 (medium shadings). The lightest shading indicates the uptake limits on 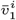. Nonnegativity constraints for fluxes (arising from reaction irreversibilities) are not shown.

The first cell (in blue color) takes up substrate *X*^1^ (its intracellular version being denoted by *S*^1^) from the medium (with maximum rate 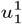), forms the biomass precursor *B*^1^ with by-product *P*^1^, and excretes the by-product (its extracellular version being denoted by *Y*^2^). Alternatively, the cell takes up substrate *Y*^1^ (its intracellular version being denoted by *A*^1^) and forms *B*^1^. As a mnemonic, *S* stands for substrate, *B* for biomass precursor, *P* for (by-)product, and *A* for alternative substrate.

The second cell (in red color) operates in a symmetric way, growing on substrate *X*^2^ with alternative substrate *Y*^2^ and (extracellular) by-product *Y*^1^.

For both cells (*i* = 1 or 2), fluxes 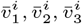 capture exchange with the medium, whereas 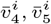 are internal fluxes. All reactions are assumed to be irreversible, that is, 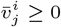.

For the first cell, we explicitly write the matrices that appear in the steady-state constraints (9a) and (9d): the (intracellular) stoichiometric matrix, the stoichiometric vector of the biomass reaction, and—above it—the corresponding extracellular stoichiometric matrix:

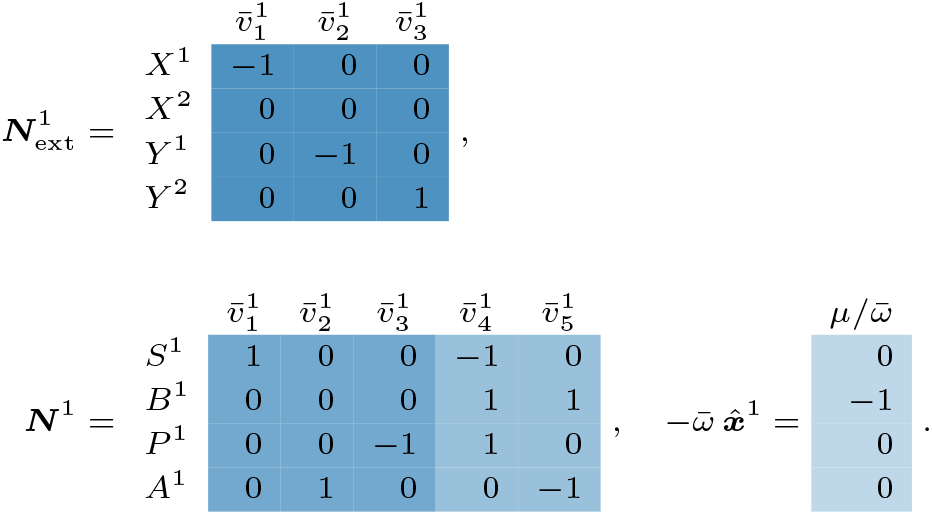

Fig. 2b summarizes the steady-state constraints of the community model, cf. (9d) and (9a), and the upper bounds for the fluxes, cf. (9b). Nonnegativity constraints for fluxes and nonnegativity and normalization constraints for microbial mass fractions, cf. (9c) and (9e), are not shown.

Altogether system (9abcd) defines a polyhedral cone. After projecting this cone to mass fractions and exchange fluxes (that is, after eliminating internal fluxes), system (11abcd) defines the composition/exchange cone 𝒞_cx_. In the example, 𝒞_cx_ is given by the equations and inequalities and by nonnegativity constraints for all variables (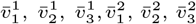, and *γ*_1_, *γ*_2_).

| constr. | $\bar{v}_1^1$ | $\bar{v}_2^1$ | $\bar{v}_3^1$ | $\gamma_1$ | $\bar{v}_1^2$ | $\bar{v}_2^2$ | $\bar{v}_3^2$ | $\gamma_2$ | sign |
| --- | --- | --- | --- | --- | --- | --- | --- | --- | --- |
| $Y^1$ | 0 | -1 | 0 | 0 | 0 | 0 | 1 | 0 | ≥ |
| $Y^2$ | 0 | 0 | 1 | 0 | 0 | -1 | 0 | 0 | ≥ |
| $S^1 + P^1$ | 1 | 0 | -1 | 0 | 0 | 0 | 0 | 0 | $=$ |
| $S^1 + B^1 + A^1$ | 1 | 1 | 0 | $-\mu/\bar{\omega}$ | 0 | 0 | 0 | 0 | $=$ |
| $S^2 + P^2$ | 0 | 0 | 0 | 0 | 1 | 0 | -1 | 0 | $=$ |
| $S^2 + B^2 + A^2$ | 0 | 0 | 0 | 0 | 1 | 1 | 0 | $-\mu/\bar{\omega}$ | $=$ |
| $\bar{v}_1^1$ | 1 | 0 | 0 | $-u_1^1$ | 0 | 0 | 0 | 0 | ≤ |
| $\bar{v}_1^2$ | 0 | 0 | 0 | 0 | 1 | 0 | 0 | $-u_1^2$ | ≤ |

In particular, for the first cell, the matrices in (11a) are:

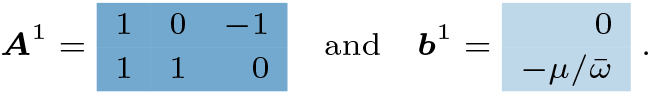

Note that the new inequalities arise as positive linear combinations of the original inequalities. For example, the constraint named *S*^1^ + *P*^1^ is obtained by adding the constraints for the intracellular metabolites *S*^1^ and *P*^1^ given in Fig. 2b. Note also that the constraints for the extracellular metabolites *X*^1^ and *X*^2^ are already covered by the nonnegativity constraints.

For symmetry reasons, we assume equal upper bounds for substrate uptake from the medium, 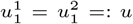, and hence 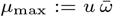 is the maximum growth rate for the two individual species. For notational simplicity, we introduce scaled fluxes 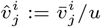 and scaled growth rate 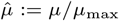, and hence 𝒞_cx_ is given by the homogeneous equations and inequalities

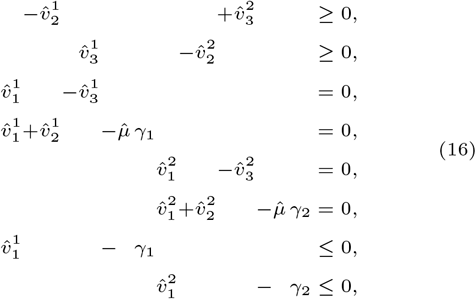

and nonnegativity constraints for all (scaled) variables.

The corresponding composition/exchange polytope 𝒫_cx_ is obtained by also considering (11e), that is, *γ*_1_ + *γ*_2_ = 1. The projection of 𝒫_cx_ to 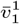 and *γ*_1_ (as a function of *µ*) is shown in Fig. 3a.

**Fig. 3.**
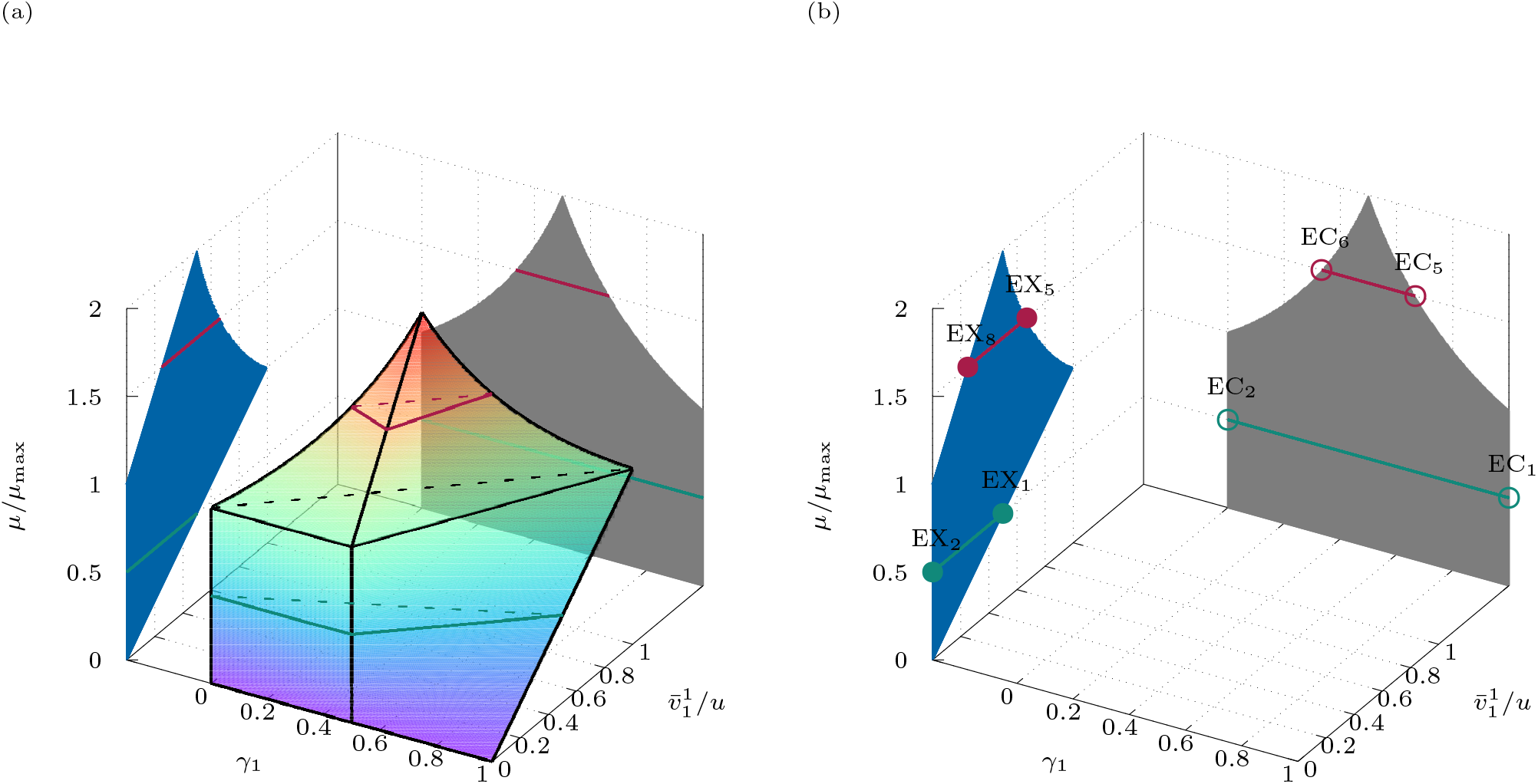
Two-species community in Fig. 2: (a) (projections of) the composition/exchange polytope as a function of growth rate, (b) (projections of) the corresponding ECs and EXs for *µ/µ*_max_ = ½ and ¾.

Finally, after projecting to mass fractions (that is, after eliminating exchange fluxes), the composition cone 𝒞_c_, is given by the homogeneous inequalities

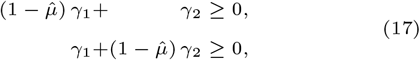

and nonnegativity constraints. As above, by also considering (11e), that is, *γ*_1_ + *γ*_2_ = 1, we obtain the composition polytope 𝒫_c_, given by the inequalities

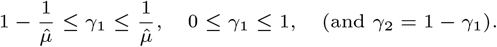

In particular, 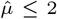 for the community (as opposed to 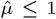 for the individual species). For fixed growth rate, feasible mass fractions *γ*_1_ lie in a certain interval: for 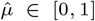, we find *γ*_1_ ∈ [0, 1]; for 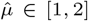, we find 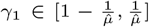. For a plot (as a function of 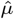), see Fig. 3b. Conversely, in terms of growth rate,

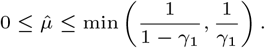

That is, for fixed mass fraction *γ*_1_, feasible growth rates 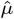 lie in a certain interval (with lower bound 0). In particular, 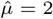 is feasible only for 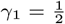.

This concludes our initial examination of Example 1, and we return to the general analysis of microbial communities. In particular, we extend the concepts of elementary flux modes and elementary conversion modes from single-species models to community models.

## Results: community elementary vectors

For the objects of polyhedral geometry (subspaces, cones, polyhedra), there is no *unique* minimal set of generators, in general. However, elementary vectors (EVs) form unique sets of *conformal* generators [39, Section 3.4] that allow to generate geometric objects *without cancellations*. For linear subspaces and s-cones (arising from linear subspaces and nonnegativity constraints) such as the flux cone [47, 48], elementary vectors are the support-minimal (SM) vectors; for general polyhedral cones such as the conversion cone [52] or the growth cone [40], they are the conformally non-decomposable (cND) vectors; finally, for polyhedra such as the flux polyhedron [27] or the growth polyhedron [40], they are the convex-conformally non-decomposable (ccND) vectors plus the cND vectors of the recession cone. For readers unfamiliar with EVs, the appendix includes an intuitive treatment in the setting of reaction networks.

Since the geometric objects 𝒫_cx_, 𝒞_cx_, 𝒫_c_, 𝒞_c_, 𝒫_x_, and 𝒞_x_ defined above are polytopes or polyhedral cones, we first recall formal definitions and results for (general) polyhedral cones and bounded polyhedra (polytopes) from [39].

### Elementary vectors

**Cones:** Let *C* be a polyhedral cone, that is, *C* = {*x* ∈ ℝ^*n*^ | *Ax* ≥ 0} for some matrix *A* ∈ ℝ^*m*×*n*^. A nonzero vector *x* ∈ *C* is *conformally non-decomposable* (cND) if, for all nonzero *x*^1^, *x*^2^ ∈ *C* with sign(*x*^1^), sign(*x*^2^) ≤ sign(*x*), the decomposition *x* = *x*^1^ + *x*^2^ implies *x*^1^ = *λx*^2^ with *λ >* 0. (If the cone is contained in an orthant, then the cND vectors coincide with the extreme rays.) In this setting, we call a vector *x* ∈ *C* an EV if it is cND. By [39, Theorem 8], every vector 0 ≠ *x* ∈ *C* is a conformal sum of EVs.

**Polytopes:** Let *P* be a polytope, in particular, *P* = {*x* ∈ ℝ^*n*^ | *Ax* ≥ *b*} for some matrix *A* ∈ ℝ^*m*×*n*^ and some vector *b* ∈ ℝ^*m*^. A vector *x* ∈ *P* is *convex-conformally non-decomposable* (ccND) if for all *x*^1^, *x*^2^ ∈ *P* with sign(*x*^1^), sign(*x*^2^) ≤ sign(*x*) and all *λ* ∈ (0, 1), the decomposition *x* = *λx*^1^ + (1 − *λ*)*x*^2^ implies *x*^1^ = *x*^2^. (If the polytope is contained in an orthant, then the ccND vectors coincide with the vertices.) In this setting, we call a vector *x* ∈ *P* an EV if it is ccND. By (a variant of) [39, Theorem 13], every vector *x* ∈ *P* is a convex-conformal sum of EVs.

Based on these definitions and results, we introduce the EVs of the polyhedral cones 𝒞_cx_, 𝒞_c_, and 𝒞_x_ and the EVs of the polytopes 𝒫_cx_, 𝒫_c_, and 𝒫_x_.

### Definition 1.

The EVs of the polyhedral cones 𝒞_cx_, 𝒞_c_, and 𝒞_x_ are their cND vectors, and the EVs of the polytopes 𝒫_cx_, 𝒫_c_, and 𝒫_x_ are their ccND vectors.

Since 𝒞_c_ and 𝒫_c_ are contained in the nonnegative orthant, we can also define their EVs as extreme rays and vertices, respectively. We name the EVs of the polytopes and cones as follows:

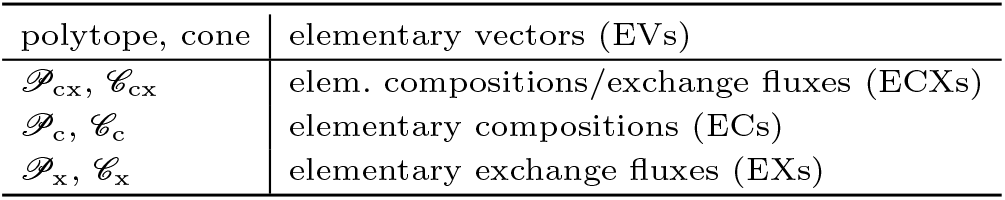

| polytope, cone | elementary vectors (EVs) |
| --- | --- |
| $\mathcal{P}_{\text{cx}}, \mathcal{C}_{\text{cx}}$ | elem. compositions/exchange fluxes (ECXs) |
| $\mathcal{P}_c, \mathcal{C}_c$ | elementary compositions (ECs) |
| $\mathcal{P}_x, \mathcal{C}_x$ | elementary exchange fluxes (EXs) |

Since corresponding cones and polytopes are isomorphic, the same holds for their EVs, and we use the same names. Indeed, the EVs of the polytopes are normalized versions of the EVs of the cones.

We explicitly state conformal decomposition results only for the composition/exchange cone and polytope. The statements for the other geometric objects are analogous.

### Theorem 3.

*Every nonzero vector x* ∈ 𝒞_cx_ *is a conformal sum of ECXs. That is, there exists a finite set E of ECXs such that*

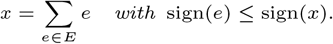

### Theorem 4.

*Every vector x* ∈ 𝒫_cx_ *is a convex, conformal sum of ECXs. That is, there exists a finite set E of normalized ECXs such that*

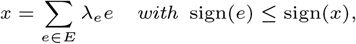

*λ*_*e*_ ≥ 0, *and* ∑_*e*∈*E*_ *λ*_*e*_ = 1.

Interestingly, ECXs and EXs are in one-to-one correspondence, if *µ >* 0.

### Proposition 5.

*If µ >* 0, *then the projection of every ECX is an EX, and every EX is the projection of exactly one ECX*.

*Proof* By projection and the linear transformations (15), respectively, elements of 𝒫_cx_ (resp. 𝒞_cx_) and elements of 𝒫_x_ (resp. 𝒞_x_) are in one-to-one correspondence. We show that every element *cx*\* ∈ 𝒞_cx_ is conformally decomposable if and only if the corresponding element *x*\* ∈ 𝒞_x_ (its projection) is conformally decomposable. For simplicity, let *cx*\* = *cx*^1^ + *cx*^2^ with *cx*^1^ ≁ *cx*^2^ and sign(*cx*^*i*^) ≤ sign(*cx*\*). Then, *x*\* = *x*^1^ + *x*^2^ with *x*^1^ ≁ *x*^2^ and sign(*x*^*i*^) ≤ sign(*x*\*) for the corresponding projections. Analogously for the other direction. □

### **Example 1** (continued)

Recall that the composition/exchange cone 𝒞_cx_ is given by Eqns. (16) and nonnegativity constraints for all (scaled) variables. The composition/exchange polytope 𝒫_cx_ is obtained by also considering the normalization *γ*_1_ + *γ*_2_ = 1. For scaled growth rate 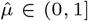, the normalized elementary compositions/exchange fluxes are given by

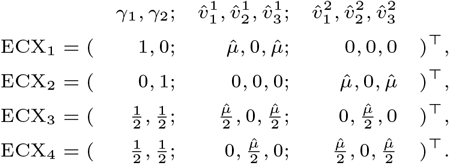

In ECX_1_ (resp. ECX_2_), only the first (resp. second) microbial species is present, whereas ECX_3_ and ECX_4_ represent true communities.

For 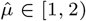, we find

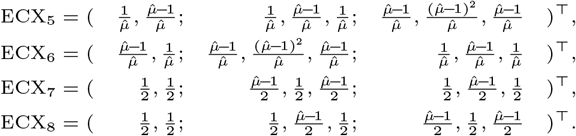

Note that ECX_5_ and ECX_6_ depend on 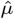 in a nonlinear way.

For 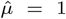 (the maximum growth rate of the individual species), the two sets of ECXs agree. For 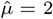 (the maximum community growth rate), there is only one ECX.

**Note** (Computation). After converting the equations and inequalities (16) into a system of equations via the introduction of slack variables, ECXs can be computed for fixed growth rate using efmtool [50]. For symbolic computation, where growth rate is treated as a parameter, we employ the SageMath packages elementary vectors and sign vectors [1, 2]; see also [3].

Further projection yields elementary compositions. For 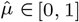, elementary compositions are given by

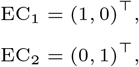

and for 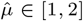, we find

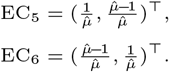

The projections of ECX_3_, ECX_4_ and ECX_7_, ECX_8_ are not ECs, but lie between EC_1_ and EC_2_ resp. EC_5_ and EC_6_. See Fig. 3b.

### Ecological significance

Elementary compositions/exchange fluxes represent *minimal modes of interaction* between members of microbial communities. For Example 1, we illustrate the ECXs in Table 2: ECX_2_ and ECX_4_ for “low” growth rate 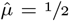 and ECX_2_ and ECX_4_ for “high” growth rate 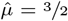. For an illustration of all ECXs, see Supplementary Table S1.

**Table 2.**
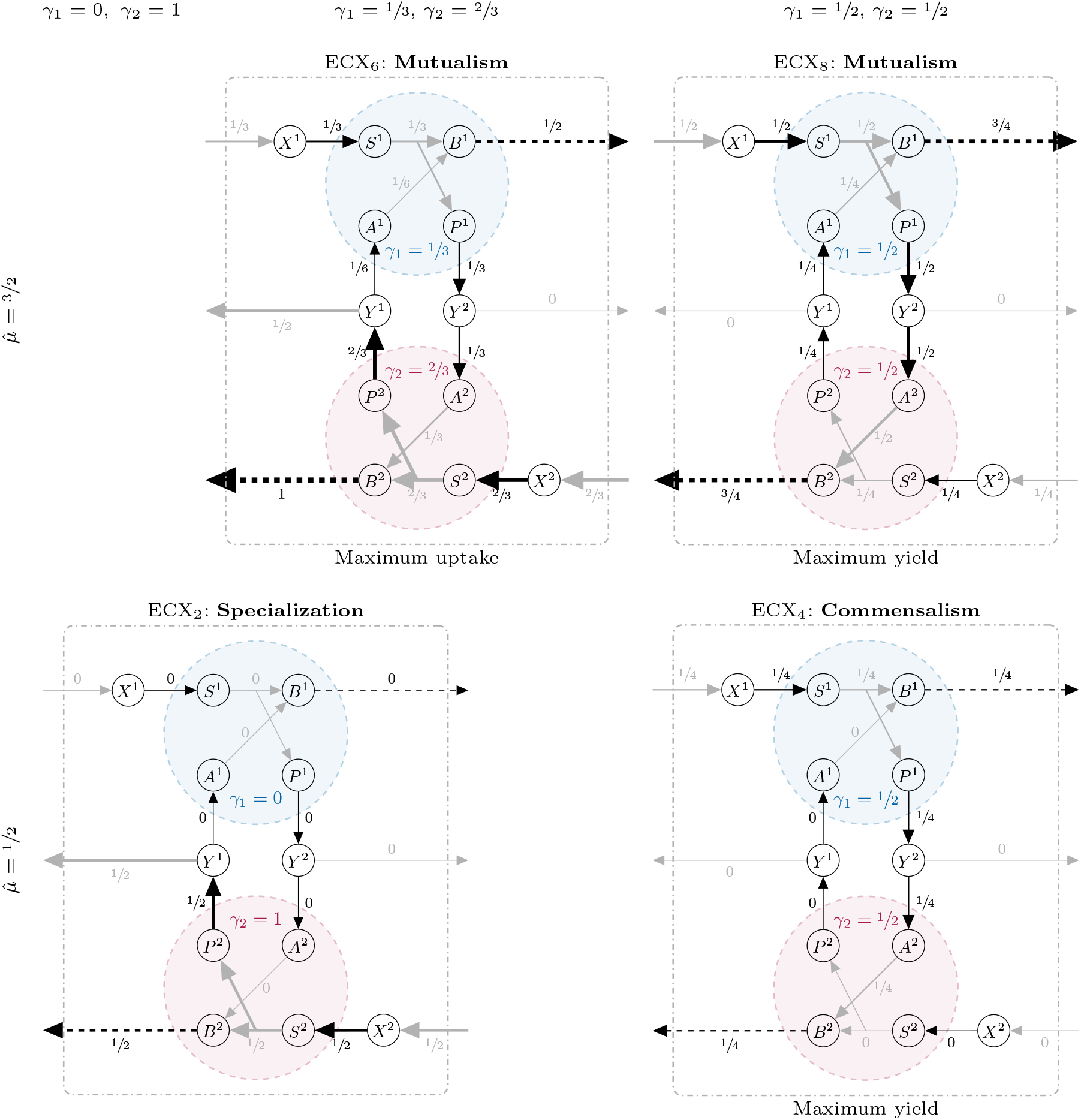
ECXs for the two-species community in Fig. 2. We show ECX_2_ and ECX_4_ for the “low” (scaled) growth rate 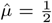 and ECX_6_ and ECX_8_ for “high” 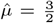. (The other ECXs are obtained by exchanging the roles of the two species.) Exchange fluxes (defining the ECXs) are shown in black, while all other fluxes are shown in gray. ECX_2_ and ECX_4_ represent specialist and commensalist minimal strategies, respectively, whereas ECX_6_ and ECX_8_ are mutualistic strategies.

**Table 3.**
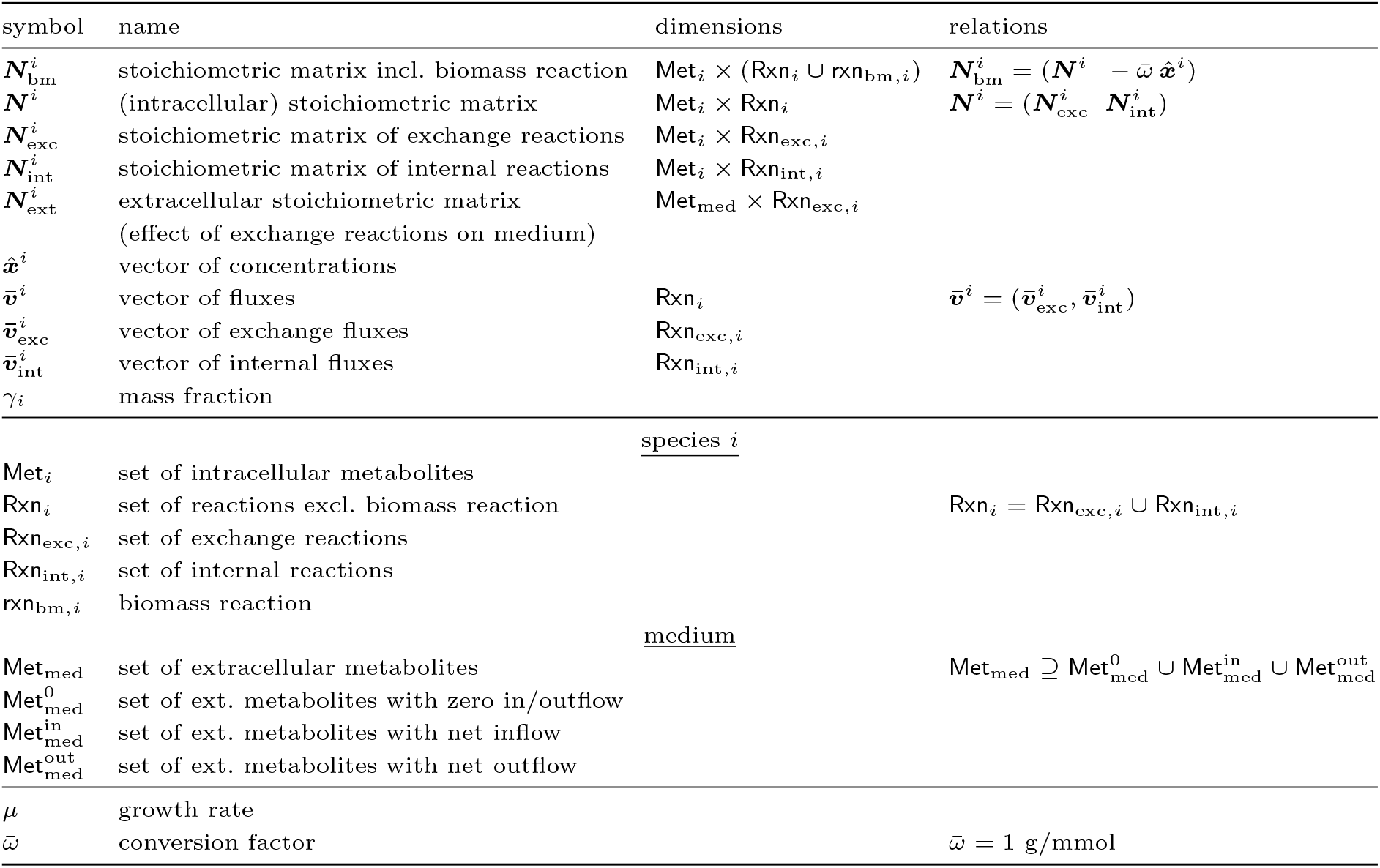
Relevant mathematical objects in the community model (9).

For generic growth rate, we find the following behaviors:

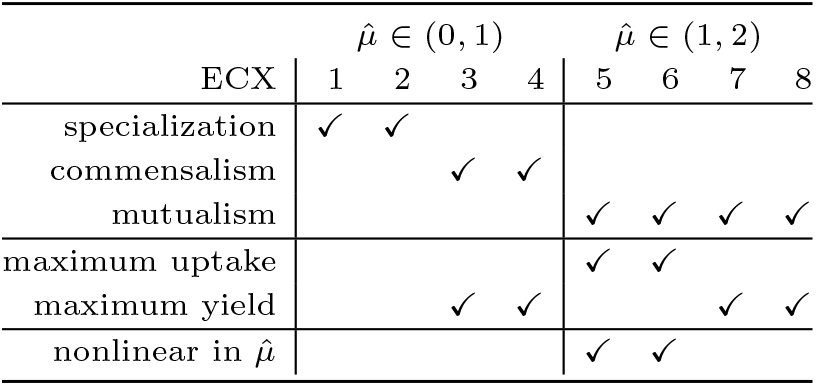

| ECX | $\hat{\mu} \in (0, 1)$ | | | | $\hat{\mu} \in (1, 2)$ | | | |
| --- | --- | --- | --- | --- | --- | --- | --- | --- |
|  | 1 | 2 | 3 | 4 | 5 | 6 | 7 | 8 |
| specialization | ✓ | ✓ |  |  |  |  |  |  |
| commensalism |  |  | ✓ | ✓ |  |  |  |  |
| mutualism |  |  |  |  | ✓ | ✓ | ✓ | ✓ |
| maximum uptake |  |  |  |  | ✓ | ✓ |  |  |
| maximum yield |  |  | ✓ | ✓ |  |  | ✓ | ✓ |
| nonlinear in $\hat{\mu}$ | | | | | ✓ | ✓ | | |

The simplest cases, ECX_1_ and ECX_2_, are paradigms for *specialization*. A species grows on its respective substrate and does not use the alternative substrate produced by the other species. *Commensalism* is represented by ECX_3_ and ECX_4_. A species growths on its substrate and produces the alternative substrate used by the other species. The most complex cases, ECX_5_, ECX_6_, ECX_7_, and ECX_8_, are paradigms of *mutualism*. Both species grow on their substrates and, additionally, use the alternative substrates produced by the respective other species. Only in this way, “high” growth rates 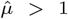 are possible. For “low” growth rates 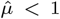, mutualism can arise from commensalism, in the sense that any convex combination of ECX_3_ and ECX_4_ exhibits mutual exchange of alternative substrates between the two species.

The ECXs can be categorized also in another way. In ECX_1_ and ECX_2_, neither substrate uptake occurs at maximum rate, nor (community) yield is maximum. (The by-product is not used as an alternative substrate and has an outflow.) On the one hand, in ECX_3_ and ECX_4_ as well in ECX_7_ and ECX_8_, no alternative substrate has an outflow, that is, the yield is maximum. On the other hand, in ECX_5_ and ECX_6_, the uptake of both substrates occurs at maximum rate. However, maximum uptake and maximum yield cannot occur at the same time, except in the case of maximum growth rate, for which there is only one ECX. To see this, recall that every element of the composition/exchange polytope is a convex, conformal sum of normalized ECXs (Theorem 4). Hence, a proper convex sum (with all coefficients nonzero) of, for example, ECX_6_ (maximum uptake) and ECX_8_ (maximum yield, but not maximum uptake) cannot exhibit maximum uptake. The same argument applies to maximum yield.

## Further examples

### Example 2

In the two-species community studied in Example 1, the projection to mass-fraction(s) and growth-rate is concave, see Figure 3b. The nonlinear part (for scaled growth rate greater than one) arises from mutualism. As we show below, cooperation can also result in convex areas.

To this end, we adapt the two-species community studied in Example 1, see Figure 4: Here, metabolite *Y*^2^ has no outflow, but species 2 can take up *Y*^2^ and, using *S*^2^, convert it to *W*^2^, which has an outflow. Moreover, the synthesis of the biomass precursor *B*^2^ and the by-product *P*^2^ are no longer coupled, but have their own (competing) synthesis reactions. In biological terms, *Y*^2^ can be interpreted as an inhibitory product, like ethanol in yeast. Species 2 can detoxify *Y*^2^, but must choose between detoxification, biomass production, and the synthesis of the product *P*^2^ which serves as an alternative substrate for species 1. Depending on the stoichiometry of biomass production and alternative substrate synthesis by species 2, the projection to mass-fraction(s) and growth-rate can be convex, linear, or concave; see Supplementary Fig. S1.

**Fig. 4.**
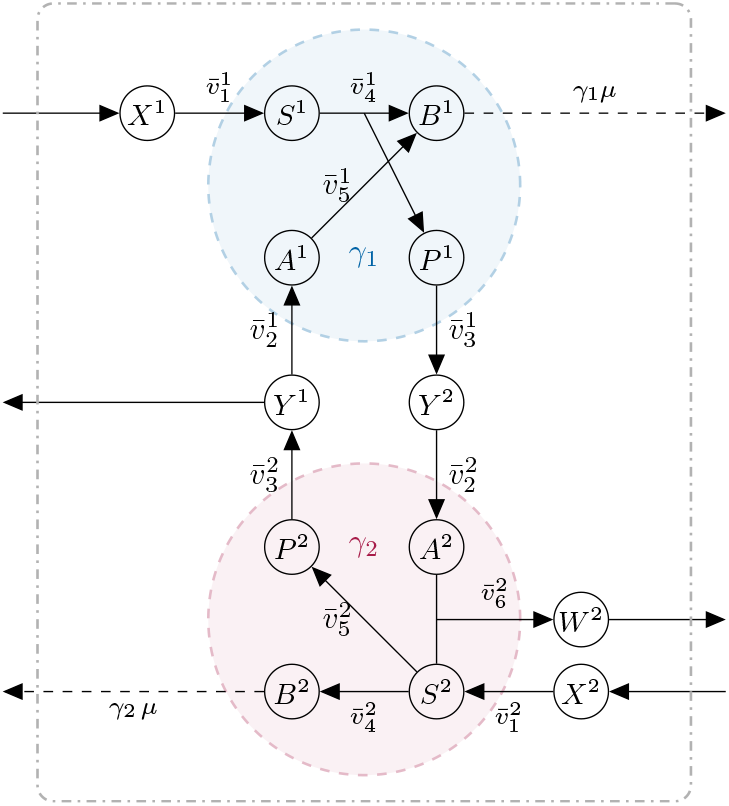
Two-species community with detoxification.

### Genome scale

#### Example 3

We consider the co-culture of *Methanobrevibacter smithii* and *Bacteroides thetaiotaomicron*, two gut microbes known to form a stable community in vitro [6, 14]. In rich medium, *M. smithii* grows slower than *B. thetaiotaomicron*, but increases its growth rate in co-culture, primarily due to hydrogen cross-feeding from *B. thetaiotaomicron*, which *M. smithii* uses for methanogenesis. Conversely, *M. smithii* also enhances the growth of *B. thetaiotaomicron*, though to a lesser extent. Beyond hydrogen, cross-feeding likely involves fermentation products, amino acids, and vitamins, most of which are suspected to be provided by *B. thetaiotaomicron*. Further, the growth of *B. thetaiotaomicron* is inhibited by some of its products, leading to a mutual benefit when these are taken up by *M. smithii* [6].

The co-culture experiments in [14] were performed in the (complex) MS medium. For our analysis of community interactions, we instead predict a *minimal* medium using the genome-scale metabolic models from [14], analogous to previous work on minimal medium design for individual organisms [17, 49, 41]. To ensure the presence of both members, we set their abundances to 1/2 in the community model (9) and use COBRApy to identify a minimal medium, based on MS medium. Specifically, we restrict the fluxes of the transport reactions to physiological ranges. As no experimental data are available, these bounds are set to ±10.0 mmol g^−1^ h^−1^, and irreversibility constraints are preserved. Inflow to the medium is bounded by 1000.0 mmol g^−1^ h^−1^, and outflow is allowed for all metabolites. The resulting minimal medium is predicted to support co-culture growth and can be prepared for in vitro testing.

In the minimal medium, *B. thetaiotaomicron* grows at 0.775 h^−1^, while *M. smithii* fails to grow in monoculture. Single knock-out analysis of MS medium metabolites identifies essential components for *M. smithii* growth that are absent from the community’s minimal medium. However, supplementing these individually does not restore growth. A combination of at least four metabolites is required: L-Proline, L-Threonine, Coenzyme A or Pantothenate, and one of 18 interchangeable compounds (mainly amino acids, fermentation products, and sugars); see Supplementary Note S1. Flux variability analysis (FVA) confirms that *B. thetaiotaomicron* supplies all required and most interchangeable metabolites. The specific metabolites exchanged (and the direction of exchange) vary with community composition, indicating the co-culture’s metabolic flexibility; see Supplementary Figure S2.

Fig. 5 shows a projection of the composition polytope (to the mass fraction of *M. smithii*) as a function of growth rate. Indeed, growth rate is varied from zero to its maximum in steps of 0.001, and FVA is used to minimize and maximize mass fractions. Recall that mass fractions can be interpreted as fluxes, and the software package PyCoMo [44] allows to apply standard CBM approaches like FVA. In this way, the computation of ECs is feasible even for genome-scale metabolic models of small communities.The maximum growth rate of 0.851 h^−1^ is reached at a community composition of 47.9% *M. smithii* (and 52.1% *B. thetaiotaomicron*). This exceeds the growth rates of either monoculture (0.0 h^−1^ for *M. smithii* and 0.755 h^−1^ for *B. thetaiotaomicron*). As in the other examples, the dependence on growth rate is non-linear.

**Fig. 5.**
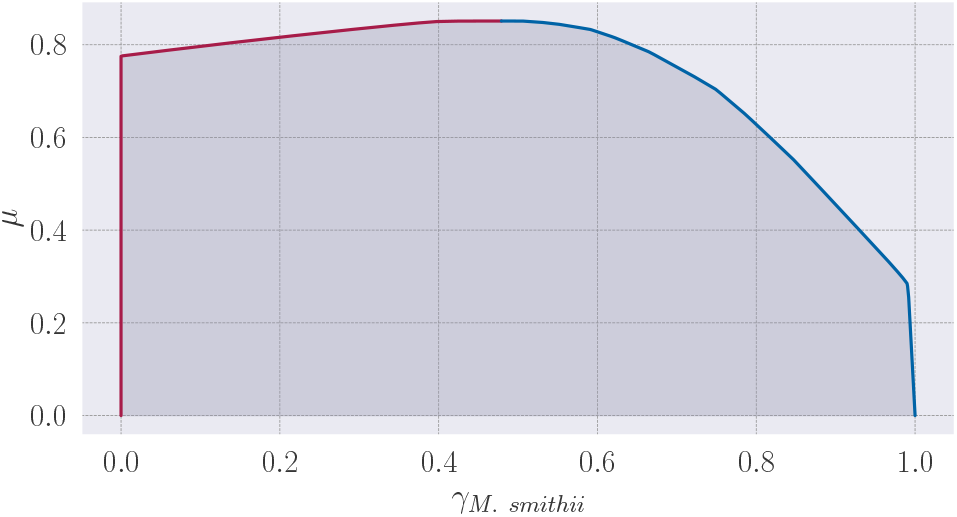
Co-culture between *M. smithii* and *B. thetaiotaomicron* on minimal medium: projection of the composition polytope (to the mass fraction of *M. smithii*) as a function of growth rate. For each growth rate, there are two elementary communities (ECs), which lie on the red and blue lines.

The mutualistic interaction arises from extensive metabolite cross-feeding. Beyond the well-established transfer of hydrogen from *B. thetaiotaomicron* to *M. smithii*, a wide range of compounds—including amino acids, sugars, nucleosides, fermentation products, and cofactors—are exchanged, consistent with prior hypotheses [6]. While most of these are supplied by *B. thetaiotaomicron*, some are produced by *M. smithii*, and for others, bidirectional exchange is possible at maximum growth; see supplement.

### Methods

Genome-scale metabolic models of *M. smithii* and *B. thetaiotaomicron* were recently reconstructed in [14], based on public genome sequences (ABYW00000000.1 and NC 004663.1), using gapseq [53]. Construction and analysis of the community metabolic model were performed using PyCoMo [44] and COBRApy [15]. A minimal medium was computed using COBRApy’s `minimal_medium` function.

Code for reproducing the results of the two-species community with detoxification (Example 2) and the co-culture of *M. smithii* and *B. thetaiotaomicron* (Example 3) is available at https://doi.org/10.5281/zenodo.17714624 [43]. Code for computing the ECXs of Example 1 using PyCoMo and efmtool [50] is available at https://doi.org/10.5281/zenodo.16744395 [38].

## Discussion

Constraint-based metabolic modeling (CBM), through techniques such as flux balance analysis (FBA) and elementary flux mode (EFM) analysis, has significantly advanced foundational research in microbiology and enabled practical applications in metabolic engineering. Central to this approach is the geometric representation of all feasible metabolic states a cell can achieve under defined conditions, known as the flux cone (or flux polyhedron in the presence of inhomogeneous constraints). This space is generated by the EFMs (or elementary flux vectors (EFVs) [27]), which represent minimal metabolic pathways and serve as fundamental building blocks of metabolism. As such, they provide a mechanistic understanding of an organism’s metabolic capabilities, independent of any optimization objective.

Despite the success of CBM in modeling individual organisms, extending it to microbial communities remains challenging and an active area of research. Optimization-based approaches have been adapted to community settings [26, 24, 29, 8, 30], yet a systematic and unbiased characterization of the full metabolic interactome has remained elusive. The core difficulty lies in generalizing the flux cone framework to multi-species systems, where community composition and interspecies interactions introduce additional layers of complexity. While EFMs represent the minimal metabolic pathways of single organisms, they fail to capture emergent community-level behaviors such as cross-feeding, syntrophy, or competition. To address this gap, we developed a geometric framework that identifies the minimal building blocks of community metabolism.

Building on existing CBM approaches for microbial communities, we defined a key geometric object: the composition and exchange flux polytope, which captures all feasible exchange fluxes between community members and their environment, along with the corresponding community compositions. Notably, the community polytope depends on both the external medium and the community growth rate.

A key implication is that methods based on the single-species flux cone can be adapted to microbial communities. Notably, the concept of EFMs extends to communities as elementary compositions/exchange fluxes (ECXs). These elementary vectors represent minimal interspecies interactions and serve as fundamental building blocks of community metabolism.

Interestingly, elementary community vectors also admit a direct ecological interpretation: each corresponds to a fundamental interaction type—specialization, commensalism, or mutualism. This connection between mechanistic modeling and ecological theory suggests that complex community-level behaviors can be decomposed into a set of elementary ecological interactions.

The identification of the community polytope also holds significant potential for biotechnological applications. Building on the flux cone framework, powerful techniques such as production envelopes and minimal cut sets [28]—widely used in metabolic engineering for single organisms—can now be extended to support the rational design and control of synthetic consortia.

A major challenge in this field is finding culture conditions that promote stable community compositions. While most interactions in natural communities are non-mutualistic [19, 42], mutualism is often desirable in synthetic systems due to its potential to enhance stability [7, 35, 34].

However, selecting compatible interaction partners and tuning environmental parameters remains a non-trivial task. In this context, elementary community vectors and their ecological interpretation offer a principled approach for identifying candidate consortia and guiding medium design.

Finally, we recall that our geometric framework relies on the assumption of balanced growth, which is well-suited to controlled biotechnological settings such as production communities maintained in steady-state bioreactors. However, this assumption may be overly restrictive for natural microbial systems, where growth rates, nutrient availability, and interaction patterns vary substantially across space and time. Relaxing the balanced-growth assumption and incorporating spatial and temporal variation would not only broaden the applicability of our framework to ecological communities, but also help clarify the conceptual foundations and assumptions underlying current community modeling approaches, including MICOM [13].

### Computation

Our framework opens many promising applications, and an important next step is to address the challenges involved in computing elementary vectors. In principle, ECXs can be obtained in two ways: (i) by first converting the full community model (9) into an equation system, enumerating its EFMs, and then projecting them (to exchange fluxes and compositions); or (ii) by first obtaining the projected community model (11)—as outlined in this work, converting it into an equation system, and enumerating its “EFMs”. The latter can also be interpreted as computing the elementary conversion modes of the full community model (9). Both strategies, however, are computationally demanding and warrant further analysis. As with classical EFM enumeration, the number of elementary vectors grows combinatorially with network size, and neither route is tractable for genome-scale community models. For small communities, consisting of up to four members, ECXs can still be obtained by the first strategy, using PyCoMo [44] to construct the equation system, but the computational cost increases rapidly with each additional species. Current algorithms, therefore, restrict ECX computation to core or pathway-level models. Developing scalable methods, whether through improved projection techniques, decomposition strategies, or approximation schemes, remains a key challenge for making ECX analysis broadly applicable to genome-scale, multi-species systems.

A promising direction for mitigating the computational burden is to first compute the elementary conversion modes [51, 5] of each species, which capture the full set of overall conversions a cell can catalyze. These elementary modes could then serve as “black-box” descriptors of the community members, encoding all possible interactions. This approach is inherently modular, as the conversion modes of each species can be computed independently and in parallel, thus reducing the community problem to several smaller, decoupled subproblems. Such a decomposition has the potential to substantially improve computational tractability and enable ECX analysis for small to moderately sized communities.

## Conflict of interest

We declare we have no competing interests.

## Data accessibility

All code and models are available at https://doi.org/10.5281/zenodo.17714624 and https://doi.org/10.5281/zenodo.16744395.

## Funding

This research was funded in whole or in part by the Austrian Science Fund (FWF), grant DOIs 10.55776/P33218 and 10.55776/PAT3748324 to SM, 10.55776/COE17 Cluster of Excellence: Circular Bioengineering to JZ, and 10.55776/J4858 to DS.

For open access purposes, the authors have applied a CC BY public copyright license to any author-accepted manuscript version arising from this submission.

## Acknowledgments

We thank Steffen Klamt^[ID](https://orcid.org/0000-0003-2563-7561)^ for discussions on community modeling and ecological aspects based on a previous version of the manuscript. We thank Marcus Aichmayr^[ID](https://orcid.org/0009-0008-8362-2644)^ and Georg Regensburger^[ID](https://orcid.org/0000-0001-7735-3726)^ for support in the symbolic computation of elementary vectors.

## Appendix: elementary vectors

In the setting of reaction networks, elementary flux modes (EFMs) are the paradigmatic examples of elementary vectors. They are defined as the *support-minimal* nonzero elements of the flux cone *C* = {**v** | **Nv** = 0 and *v*_*i*_ ≥ 0 for *i* ∈ *I*}, where *I* is the set of irreversible reactions. They represent the minimal sets of reactions that can carry steady-state flux. Most importantly, every feasible flux can be written as a conformal sum (a sum without cancellations) of EFMs; in this sense, the set of EFMs forms the minimal building blocks of a metabolic network when modeled by the flux cone, that is, using only stoichiometric (and irreversibility) constraints.

When additional linear constraints such as flux bounds or enzyme capacities are considered (summarized as **Av** ≥ **b**), the feasible fluxes form a flux polyhedron. Its elementary vectors are called elementary flux vectors (EFVs). They are not support-minimal in general, but conformally non-decomposable, that is, they cannot be written as a sum of two vectors without cancellations. (The technical definition is more involved, and we omit it here to keep the treatment intuitive.)

In the simple network shown in Fig. 6a, there are three irreversible reactions (and the flux bound *v*_1_ ≤ 1/2). The flux cone shown in Fig. 6b is defined by stoichiometric and irreversibility constraints (but not by the flux bound). It has two representative EFMs: EFM_1_ = (1, 0, 1)^⊤^ uses reactions 1 and 3, whereas EFM_2_ = (0, 1, 1)^⊤^ uses reactions 2 and 3. Every element of the cone is a nonnegative sum of the two EFMs. (Here, the sum is trivially without cancellations, since all reactions are irreversible.)

**Fig. 6.**
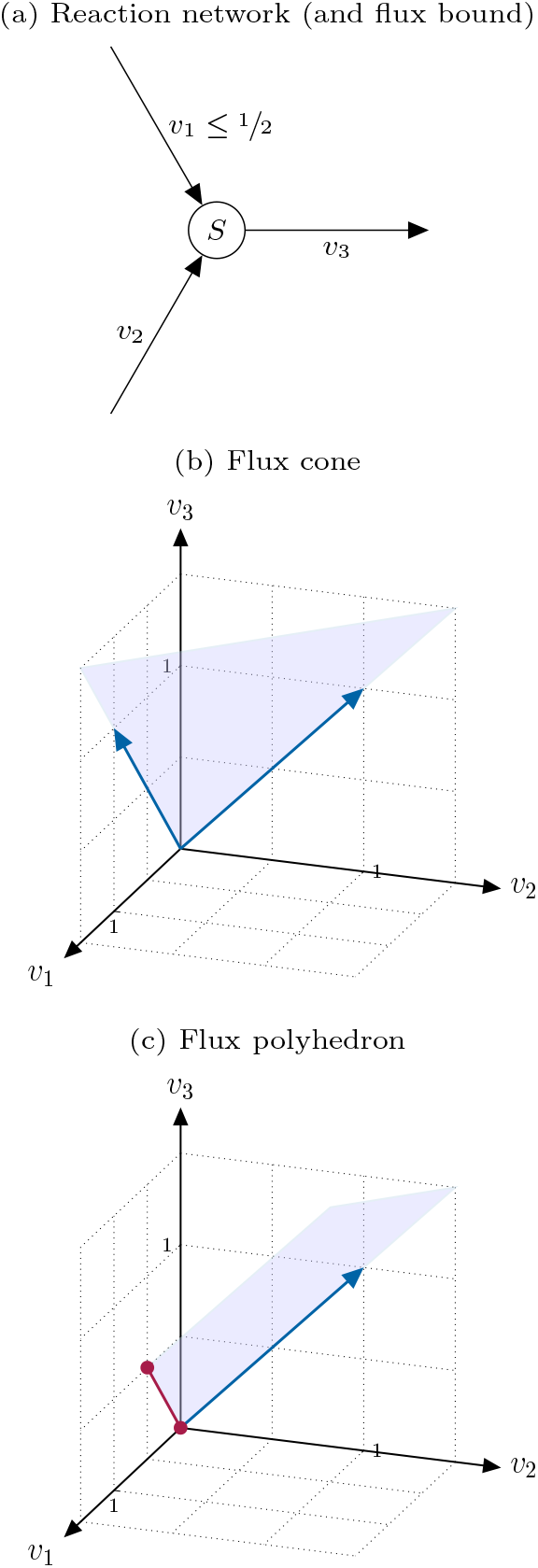
Elementary vectors for reaction networks: EFMs of the flux cone and EFVs of the flux polyhedron

The flux polyhedron shown in Fig. 6c is defined by all constraints. It has three EFVs: EFV_1_ = (1/2, 0, 1/2)^⊤^ is limited by the flux bound on reaction 1, EFV_2_ = EFM_2_ = (0, 1, 1)^⊤^ is not affected by the flux bound, and EFV_3_ = (0, 0, 0)^⊤^ represents zero flux. Every element of the polyhedron is a convex sum of EFV_1_ and EFV_3_ (red dots) plus a nonnegative multiple of EFV_2_ (blue arrow).

## Footnotes

1 To be precise, we should define 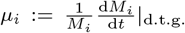, where d.t.g. denotes “due to growth”. In particular, in a flow reactor, the dynamics is determined by growth and dilution, 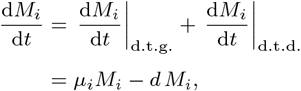 where d.t.d. denotes “due to dilution” and *d* is the dilution rate.

2 Alternatively, we may introduce the vector of mol fractions 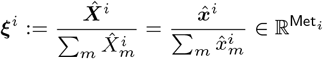 with 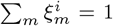. Using the mass constraint (2), we find 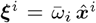 with average molar mass 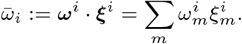 Then, the stoichiometric vector of the biomass reaction is −***ξ***^*i*^ and the biomass flux is 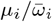.

3 Additionally/alternatively, we may consider the enzyme capacity constraint 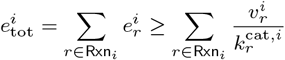 together with nonnegativity constraints for all (forward and backward) fluxes (after “reaction splitting”). All linear constraints can be summarized as ***A***^*i*^***v***^*i*^ ≥ ***b***^*i*^ with a matrix 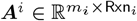 and a vector 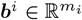.

4 More generally, we may restrict net out/inflows to given intervals. More specifically, for *j* ∈ Met_med_, we may assume Φ_*j*_ ∈ Int_*j*_ ⊆ ℝ. Then, 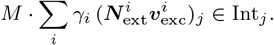 It turns out that there are 15 possible intervals, given the signs of their lower/upper bounds. Here, we state a few relevant cases. 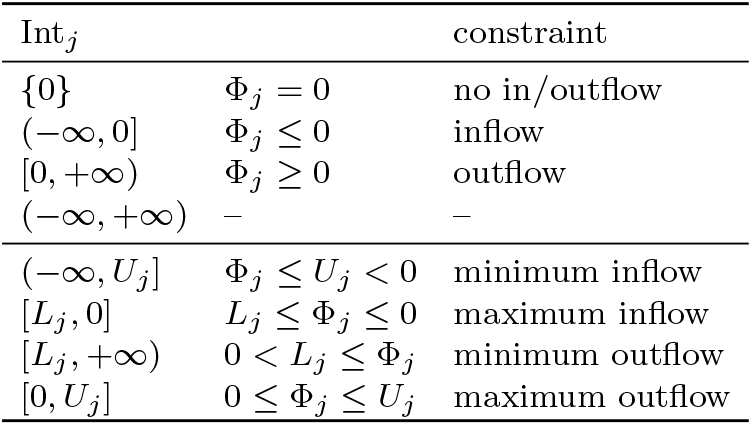 In this work, we consider only the homogeneous constraints introduced above to obtain the simplest geometric objects.

## Notes

### Competing Interest Statement

The authors have declared no competing interest.

### Summary of Updates

We have improved the presentation in the main text and updated the supplementary material accordingly.

