## Supplementary material for "Elementary vectors reveal minimal interactions in microbial communities"

#### Contents

Supplementary Figures 2
Supplementary Tables 3
Supplementary Notes 6

#### List of Figures

S1 Two-species model with detoxification 2
S2 Cross-fed metabolites in a co-culture of *M. smithii* and *B. thetaiotaomicron* 3

#### List of Tables

S1 ECXs for the two-species community 4
S2 Characteristics of the genome-scale metabolic models 5
S3 Composition of MS medium 5
S4 Compositions of the minimal media 5

### Supplementary Figures

(a) Convex: $S^2 \rightarrow {}^1\!/_2\, B^2$.

(b) Linear: $S^2 \rightarrow 1\, B^2$.

(c) Concave: $S^2 \rightarrow 2\, B^2$

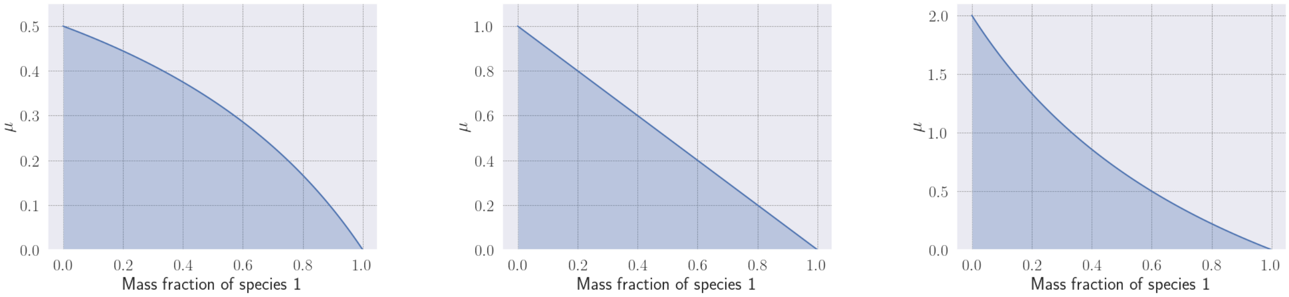


**Fig. S1.** Two-species model with detoxification (Example 2 in the main text): projections to growth rate and mass fraction of species 1. Feasible areas have convex, linear, or concave shapes depending on the stoichiometry $b$ of the internal reaction $S^2 \rightarrow b\, B^2$ of species 2. Computations were performed in COBRApy [2]; the mass fraction of species 1 was varied from 0 to 1 in steps of 0.01, and corresponding minimum and maximum growth rates were computed.

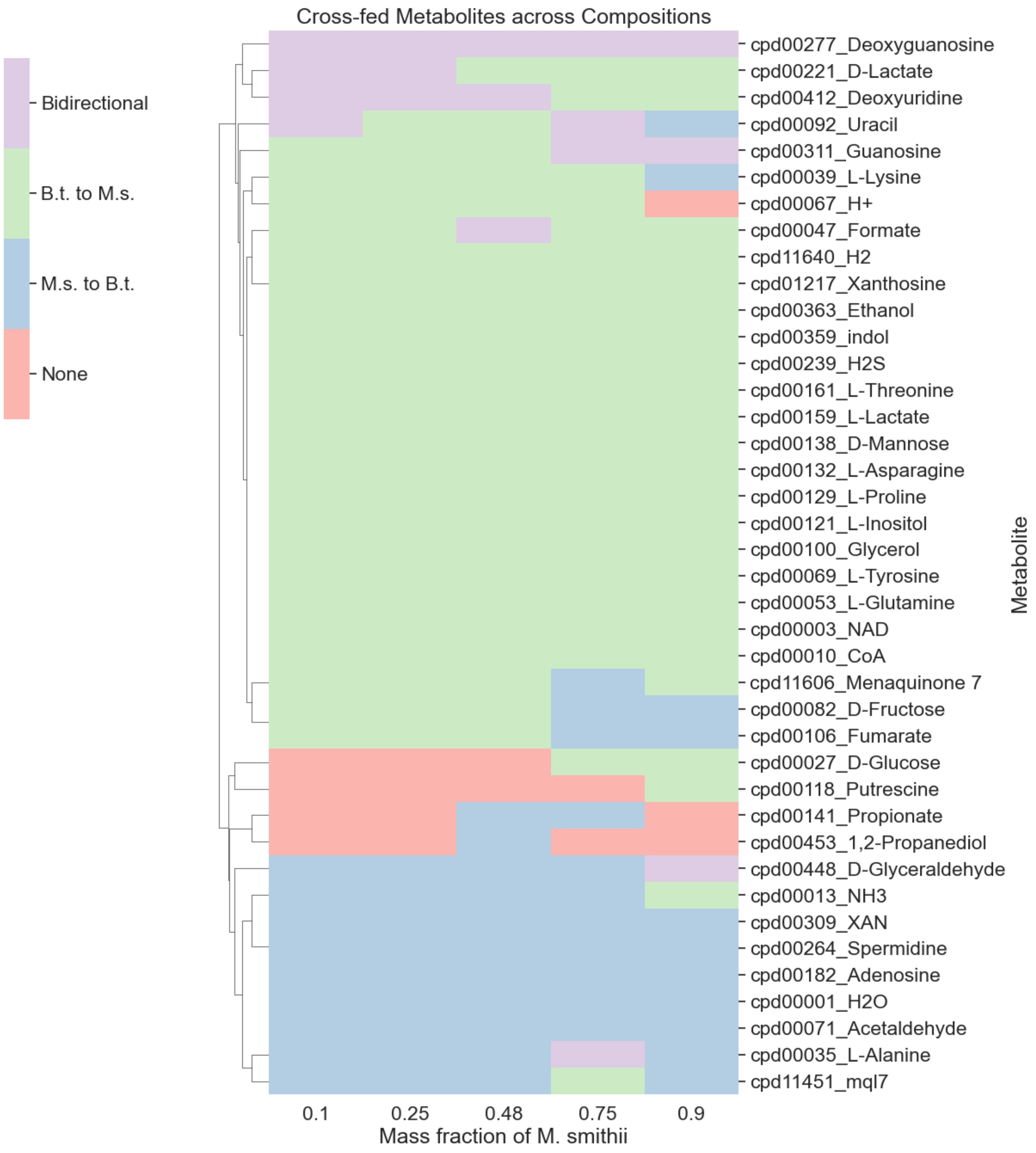


**Fig. S2.** Cross-fed metabolites in a co-culture of *M. smithii* and *B. thetaiotaomicron* (Example 3 in the main text) across five mass fractions of *M. smithii*. (Maximum growth occurs at mass fraction 0.48.) Cross-feeding was inferred using FVA performed at the maximum growth rate for each mass fraction: a metabolite is considered cross-fed if the flux ranges allow excretion by one species and uptake by the other. Metabolites present in the medium or not cross-fed at any sampling point are excluded.

#### Supplementary Tables

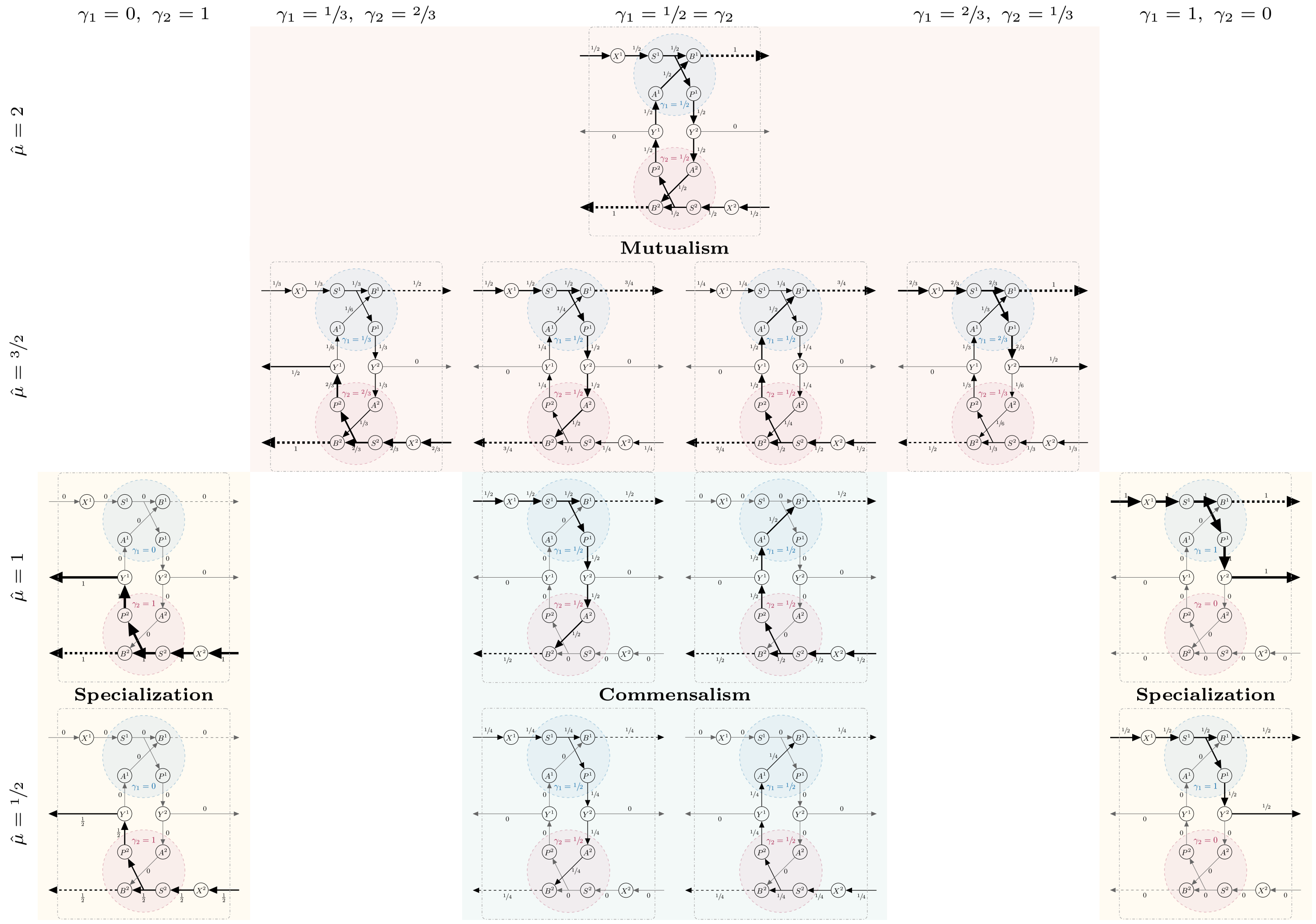


**Table S1.** ECXs for the two-species community in Fig. 2 in the main text for (scaled) growth rates $\hat{\mu} = 1/2, 1, 3/2, 2$. At maximum individual growth rate $\hat{\mu} = 1$, there is a bifurcation from specialist/commensal minimal strategies to mutualistic strategies. At maximum community growth rate $\hat{\mu} = 2$, the four ECXs collapse into one.

| | *B. thetaiotaomicron* | *M. smithii* |
|---|---|---|
| Reactions | 2087 | 1558 |
| Metabolites | 1672 | 1481 |
| Genes | 708 | 433 |

**Table S2.** Structural characteristics of the two genome-scale metabolic models used in Example 3 in the main text. See [1].

| **MS medium without additives** | | | |
|---|---|---|---|
| **compound** | **name** | **compound** | **name** |
| cpd00001 | H2O | cpd04097 | Lead |
| cpd09693 | Ba2+ | cpd00030 | Mn2+ |
| cpd01012 | Cadmium | cpd00149 | Cobalt |
| cpd09695 | Sr2+ | cpd00034 | Zn2+ |
| cpd11744 | Vanadate | cpd00058 | Cu2+ |
| cpd24346 | Sn++ | cpd23913 | aluminum chloride |
| cpd00131 | Molybdenum | cpd09225 | Boric acid |
| cpd00128 | Adenine | cpd11574 | Molybdate |
| cpd00207 | Guanine | cpd15574 | tungstate |
| cpd00307 | Cytosine | cpd03396 | Selenate |
| cpd00092 | Uracil | cpd00104 | Biotin |
| cpd90020 | cellulose | cpd00393 | Folate |
| cpd00138 | D-Mannose | cpd00263 | Pyridoxol |
| cpd01079 | Selenium | cpd00305 | Thiamin |
| cpd00059 | L-Ascorbate | cpd00220 | Riboflavin |
| cpd00098 | Choline | cpd00218 | Niacin |
| cpd00035 | L-Alanine | cpd00644 | Pantothenic acid |
| cpd19181 | Aspartate | cpd00423 | Cob(II)alamin |
| cpd00132 | L-Asparagine | cpd00443 | 4-Aminobenzoate |
| cpd00023 | L-Glutamate | cpd14958 | (R)-Lipoic acid |
| cpd00253 | Glutamine | cpd00242 | H2CO3 |
| cpd00033 | Glycine | cpd00254 | Mg |
| cpd00119 | L-Histidine | cpd00205 | K+ |
| cpd00322 | L-Isoleucine | cpd00009 | Phosphate |
| cpd00107 | L-Leucine | cpd00063 | Ca2+ |
| cpd00549 | D-Lysine | cpd00099 | Cl- |
| cpd00060 | L-Methionine | cpd00013 | Ammonium |
| cpd00066 | L-Phenylalanine | cpd00244 | Nickel |
| cpd00129 | L-Proline | cpd10515 | Fe2+ |
| cpd00054 | L-Serine | cpd10516 | Fe3+ |
| cpd00161 | L-Threonine | cpd00048 | Sulfate |
| cpd00065 | L-Tryptophan | cpd08053 | Resazurin |
| cpd00069 | L-Tyrosine | cpd00084 | L-Cysteine |
| cpd00156 | L-Valine | cpd00239 | H2S |
| cpd00051 | L-Arginine | cpd00971 | Sodium |
| cpd20624 | Chromium (III) | cpd00029 | Acetate |
| **Additives to support growth of *B. thetaiotaomicron*** | | | |
| cpd01401 | Vitamin K1 | cpd04145 | Hemine |

**Table S3.** Composition of MS medium plus additives [1] to support growth of *B. thetaiotaomicron*.

| **Minimal media: common compounds** | |
|---|---|
| **compound** | **name** |
| cpd00009 | Phosphate |
| cpd00048 | Sulfate |
| cpd00051 | L-Arginine |
| cpd00060 | L-Methionine |
| cpd00084 | L-Cysteine |
| cpd00119 | L-Histidine |
| cpd00149 | Cobalt |
| cpd00244 | Nickel |
| cpd00393 | Folate |

| ***M. smithii* mono-culture** | | **co-culture** | |
|---|---|---|---|
| **compound** | **name** | **compound** | **name** |
| cpd00065 | L-Tryptophan | cpd00030 | Mn2+ |
| cpd00066 | L-Phenylalanine | cpd00034 | Zn2+ |
| cpd00069 | L-Tyrosine | cpd00063 | Ca2+ |
| cpd00092 | Uracil | cpd00058 | Cu2+ |
| cpd00098 | Choline | cpd00099 | Cl- |
| cpd00129 | L-Proline | cpd00205 | K+ |
| cpd00132 | L-Asparagine | cpd00220 | Riboflavin |
| cpd00138 | D-Mannose | cpd00254 | Mg |
| cpd00156 | L-Valine | cpd10515 | Fe2+ |
| cpd00161 | L-Threonine | cpd10516 | Fe3+ |
| cpd00322 | L-Isoleucine | cpd90020 | cellulose |
| cpd00644 | Pantothenic acid | | |

**Table S4.** Compositions of the minimal media predicted for *M. smithii* mono-culture and co-culture with *B. thetaiotaomicron*. Common components on top, followed by compounds unique to the mono-culture medium (left) and co-culture medium (right). Predictions computed with COBRApy `minimal_medium` function [2] based on the additive-supplemented MS medium (Supplementary Table S3).

#### Supplementary Notes

**Note S1.** Supplementation of community minimal medium for rescuing growth of M. smithii

*M. smithii* has five (single knock out) essential metabolites:

| | |
|---|---|
| Cpd00129 | L-Proline |
| Cpd00149 | Co2+ (Cobalt 2+) |
| Cpd00161 | L-threonine |
| Cpd00244 | Ni2+ (Nickel 2+) |
| Cpd00393 | Folate |

Of those single KO essential metabolites, Cobalt, Nickel and Folate are included in the community minimal medium, but not L-proline and L-threonine.
Supplementing the community minimal medium with the two amino acids is, however, not sufficient for rescuing growth.
At least 4 metabolites are required as supplements to rescue the growth of *M. smithii*. When sampling all exchange reactions of the model, a total of 36 combinations of 4 metabolites is capable of rescuing growth (growth rate $>$ 0.001). Interestingly, these combinations vary mostly in only a single metabolite and can be formed as follows:

| **Missing essentials (all required)** | |
|---|---|
| Cpd00129 | L-Proline |
| Cpd00161 | L-threonine |
| **Form of Vitamin B5 (one required)** | |
| Cpd00010 | CoA (Coenzyme A) |
| Cpd00644 | Pantothenate |
| **Additional substrate (one required)** | |
| Cpd00001 | Water |
| Cpd00047 | Formate |
| Cpd01861 | Acenaphthoquinone |
| Cpd00453 | 1,2-Propanediol |
| Cpd00363 | Ethanol |
| Cpd00055 | Formaldehyde |
| Cpd00448 | D-Glyceraldehyde |
| Cpd11640 | H2 |
| Cpd00116 | Methanol |
| Cpd00138 | D-Mannose |
| Cpd00027 | D-Glucose |
| Cpd00082 | D-Fructose |
| Cpd00159 | L-Lactate |
| Cpd00221 | D-Lactate |
| Cpd00100 | Glycerol |
| Cpd00105 | D-Ribose |
| Cpd00412 | Deoxyuridine |
| Cpd01217 | Xanthosine |

Highest growth rates could be achieved by D-Fructose, D-Mannose and D-Glucose (0.61, 0.58, 0.45). H2 could only yield a growth rate of 0.18.
Of those 18, the following are not provided by *B. thetaiotaomicron*:

| | |
|---|---|
| Cpd01861 | Acenaphthoquinone |
| Cpd00001 | Water |
| Cpd00453 | 1,2-Propanediol |
| Cpd00055 | Formaldehyde |
| Cpd00116 | Methanol |
| Cpd00105 | D-Ribose |

Of those metabolites, only water is cross-fed, but from *M. smithii* to *B. thetaiotaomicron* (see supplementary figure S2). Further, while D-Glyceraldehyde can be provided by *B. thetaiotaomicron*, it appears to be mostly consuming it, according to the FVA analyses.
This means, that all three essential (only CoA, not Pantothenate) and 12 of 18 (66.7%) potentially required metabolites are provided by *B. thetaiotaomicron* during co-culture. This presents a high degree of metabolic flexibility, as can also be seen in the analysis of cross-fed metabolites across different compositions.
